# Genome-Wide Mapping of RNA-Protein Associations via Sequencing

**DOI:** 10.1101/2024.09.04.611288

**Authors:** Zhijie Qi, Shuanghong Xue, Junchen Chen, Wenxin Zhao, Kara Johnson, Xingzhao Wen, John Lalith Charles Richard, Sheng Zhong

## Abstract

RNA-protein interactions are crucial for regulating gene expression and cellular functions, with their dysregulation potentially impacting disease progression. Systematically mapping these interactions is resource-intensive due to the vast number of potential RNA and protein interactions. Here, we introduce PRIM-seq (<u>P</u>rotein-<u>R</u>NA Interaction <u>M</u>apping by sequencing), a method for the concurrent *de novo* identification of RNA-binding proteins (RBPs) and the elucidation of their associated RNAs. PRIM-seq works by converting each RNA-protein pair into a unique chimeric DNA sequence, which is then decoded through DNA sequencing. Applied to two human cell types, PRIM-seq generated a comprehensive human RNA-protein association network (HuRPA), consisting of more than 350,000 RNA-proteins pairs involving approximately 7,000 RNAs and 11,000 proteins. The data revealed an enrichment of previously reported RBPs and RNA-protein interactions within HuRPA. We also identified LINC00339 as a protein-associating non-coding RNA and PHGDH as an RNA-associating protein. Notably, PHGDH interacts with BECN1 and ATF4 mRNAs, suppressing their protein expression and consequently inhibiting autophagy, apoptosis, and neurite outgrowth while promoting cell proliferation. PRIM-seq offers a powerful tool for discovering RBPs and RNA-protein associations, contributing to more comprehensive functional genome annotations.

## Introduction

RNA-protein associations are fundamental to the intricate and dynamic processes of cellular life. These interactions underpin a multitude of biological functions, from gene expression regulation to the maintenance of cellular structure. At the molecular level, the interplay between RNA and proteins orchestrates the synthesis, processing, and regulation of RNA molecules, influencing almost every aspect of cellular physiology ^1–4^.

One of the most critical roles of RNA-protein associations is in the regulation of gene expression. RNA-binding proteins (RBPs) interact with various RNA species to control their stability, localization, and translation ^5–9^. These interactions can dictate the fate of an RNA molecule, determining whether it is translated into a protein, degraded, or sequestered in specific cellular compartments ^10^. Beyond gene expression, RNA-protein associations play a significant role in maintaining RNA structure and integrity, preventing misfolding or aggregation ^11,12^. Furthermore, RNA-protein associations are crucial in RNA transport, splicing, and editing, highlighting their versatile roles in post-transcriptional regulation ^13,14^. Dysregulation of RNA-protein associations is often associated with diseases, including cancer, neurodegenerative disorders, and viral infections, making them a focal point for therapeutic interventions ^15–17^.

Significant advances in identifying RNA-protein associations are made via two technical routes, including characterization of proteins bound to an RNA of interest (RNA-centric), and examination of RNAs bound to a protein of interest (protein-centric) ^18^. RNA-centric approaches involve the purification of specific RNAs followed by the identification of co-purified proteins ^2,4,19^, including RNA interactome capture (RIC) ^20^, click chemistry-assisted RBA interactome capture (CARIC) ^21^, RNA interactome using click chemistry (RICK) ^22^, RNA affinity purification (RAP) ^23^, tandem RNA isolation procedure (TRIP) ^23,24^, peptide-nucleic-acid-assisted identification of RNA-binding proteins (PAIR) ^25^, and MS2 in vivo biotin-tagged RAP (MS2-BioTRAP) ^26^, and RNA-protein interaction detection (RaPID) ^27^. Furthermore, proximity labeling technologies attach a labeling enzyme to the RNA of interest and capture the RBPs in the vicinity ^28–30^. Additionally, CRISPR-based RNA-United Interacting System (CRUIS) ^31^, CRISPR-based RNA proximity proteomics (CBRPP) ^32^, and in-cell protein-RNA interaction (incPRINT) leverages the RNA CRISPR system to label the proteins associated with the RNA of interest ^20^.

Protein-centric approaches purify a specific RBP and identify the co-purified RNAs, including RNA-co-immunoprecipitation followed by sequencing (RIP-seq) ^33^, and crosslinking and immunoprecipitation followed by sequencing (CLIP-seq) ^34^, as well as variations of CLIP-seq including PAR-CLIP ^35^, individual-nucleotide resolution UV crosslinking and immunoprecipitation (iCLIP) ^36^, high-throughput sequencing of RNA isolated by crosslinking immunoprecipitation (HITS-CLIP) ^37^, enhanced CLIP (eCLIP) ^38,39^, BrdU-CLIP ^40^, infrared-CLIP (irCLIP) ^41^, Fusion-CLIP ^42^, GoldCLIP ^43^, and esayCLIP ^44^. Additionally, targets of RNA-binding proteins identified by editing (TRIBE) ^45^, HyperTRIBE ^46^, and targets of RBPs identified by editing induced through dimerization (TRIBE-ID) ^47^, fuse the RNA-editing enzyme ADAR with the RBP of interest to define RNA targets by RNA editing events.

Despite significant advances, identifying the entire network of RNA-protein associations remains a formidable challenge. The human genome comprises approximately 60,000 annotated genes, with around 30,000 protein-coding and 30,000 non-coding genes. Generating a comprehensive pairwise RNA-protein interaction map is daunting due to the vast number of potential interactions, estimated at 60,000 × 30,000 candidate pairs, not even accounting for splicing variants. Existing technologies are limited in capacity, typically focusing on one RNA or RBP at a time and operating on a "one-to-many" mapping scale. In response to this limitation, we propose a "many-to-many" mapping strategy with PRIM-seq (<u>P</u>rotein-<u>R</u>NA Interaction <u>M</u>apping by sequencing), which concurrently identifies RNA-associating proteins and their associated RNAs at the genome scale. PRIM-seq does not require specific reagents to target individual proteins or RNAs, allowing for the de novo identification of RNA-protein associations.

## Results

### The PRIM-seq technology

The fundamental concept behind PRIM-seq is to transform each RNA-protein association into a chimeric DNA sequence, where a part of this chimeric sequence reflects the protein and the other part of this chimeric sequence reflects the associated RNA. These chimeric DNA sequences are sequenced by high-throughput sequencing and used for decoding the RNA-protein associations. These chimeric sequences are created by two steps. First, a protein library is created where each protein is conjugated with its own mRNA (Figure 1a, Figure S1a) ^48^. Second, mRNA is converted to cDNA, creating a library of cDNA-labeled proteins. These proteins are allowed to interact with the transcriptome retrieved from the cells of interest. Protein-associated RNA is ligated with the cDNA of the associating protein, creating a cDNA-RNA chimeric sequence. These cDNA-RNA chimeric sequences are converted to DNA sequences for sequencing (Figure 1b).

**Figure 1.**
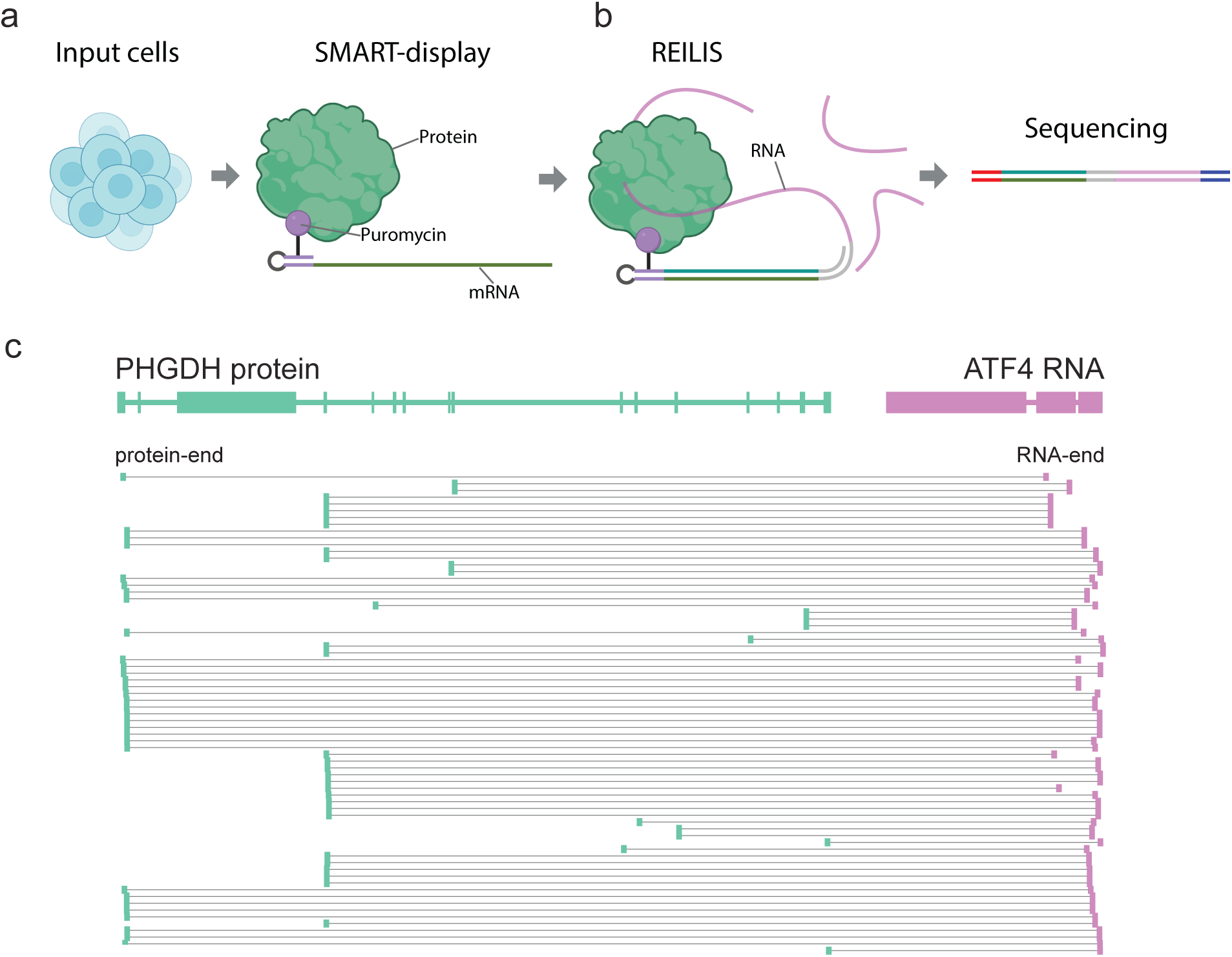
PRIM-seq experimental pipeline. (a) Step 1, SMART-display takes mRNA from input cells to produce a library of mRNA-barcoded proteins. The protein and its mRNA label are covalently linked through puromycin (b) Step 2, REILIS converts each RNA-protein pair to a chimeric DNA sequence, with a “protein-end” read originating from the protein’s mRNA label (green) and a “RNA-end” read originating from the associated RNA (purple). (c) PRIM-seq read pairs with the protein-end aligned to the PHGDH gene and the RNA-end aligned to the ATF4 gene.

PRIM-seq comprises two experimental modules and a bioinformatics module. The first experimental module utilizes the recently developed SMART-display technology, which labels tens of thousands of proteins with their own mRNA to generate a library of RNA-labeled proteins ^48^ (Figure 1a). Briefly, SMART-display extracts mRNAs from input cells, removes the polyA tails, appends a puromycin-conjugated linker to the 3’ ends of these mRNAs, and translates the mRNAs to yield proteins that are covalently linked with the mRNAs (mRNA-linker-protein) ^48^. A key advantage of SMART-display is its capability to create the entire library of mRNA-labeled proteins in a one-pot procedure, eliminating the need for RNA-, gene-, or protein-specific reactions.

The second experimental module is called REILIS (<u>Re</u>verse transcription, Incubation, <u>L</u>igation, and <u>S</u>equencing). REILIS transforms each RNA-protein pair into a chimeric sequence that includes an “RNA-end read” and a “protein-end read” representing the associating RNA and protein, respectively (Figure 1b, Figure 2, Figure S1). REILIS takes two inputs: a library of mRNA-labeled proteins generated by SMART-display and an RNA library. REILIS converts each protein’s mRNA label into a cDNA label (Figure 2a,b), incubates the protein library with an RNA library to allow for RNA-protein association (Figure 2c), ligates the RNA and the cDNA label of each RNA-protein pair into a chimeric sequence to form an RNA-linker-cDNA structure (Figure 2d), and subjects this chimeric sequence to paired-end sequencing (Figure 2e-i, Figure S1). The cDNA fraction of this chimeric sequence reflects the protein (protein-end) and the RNA fraction reflects the associated RNA (RNA-end, Figure 2i). We note that REILIS is designed to capture RNA-protein pairs from RNA-protein complexes, which are crosslinked by formaldehyde; therefore, REILIS (and PRIM-seq) is not intended exclusively for detecting direct protein-RNA binding.

**Figure 2.**
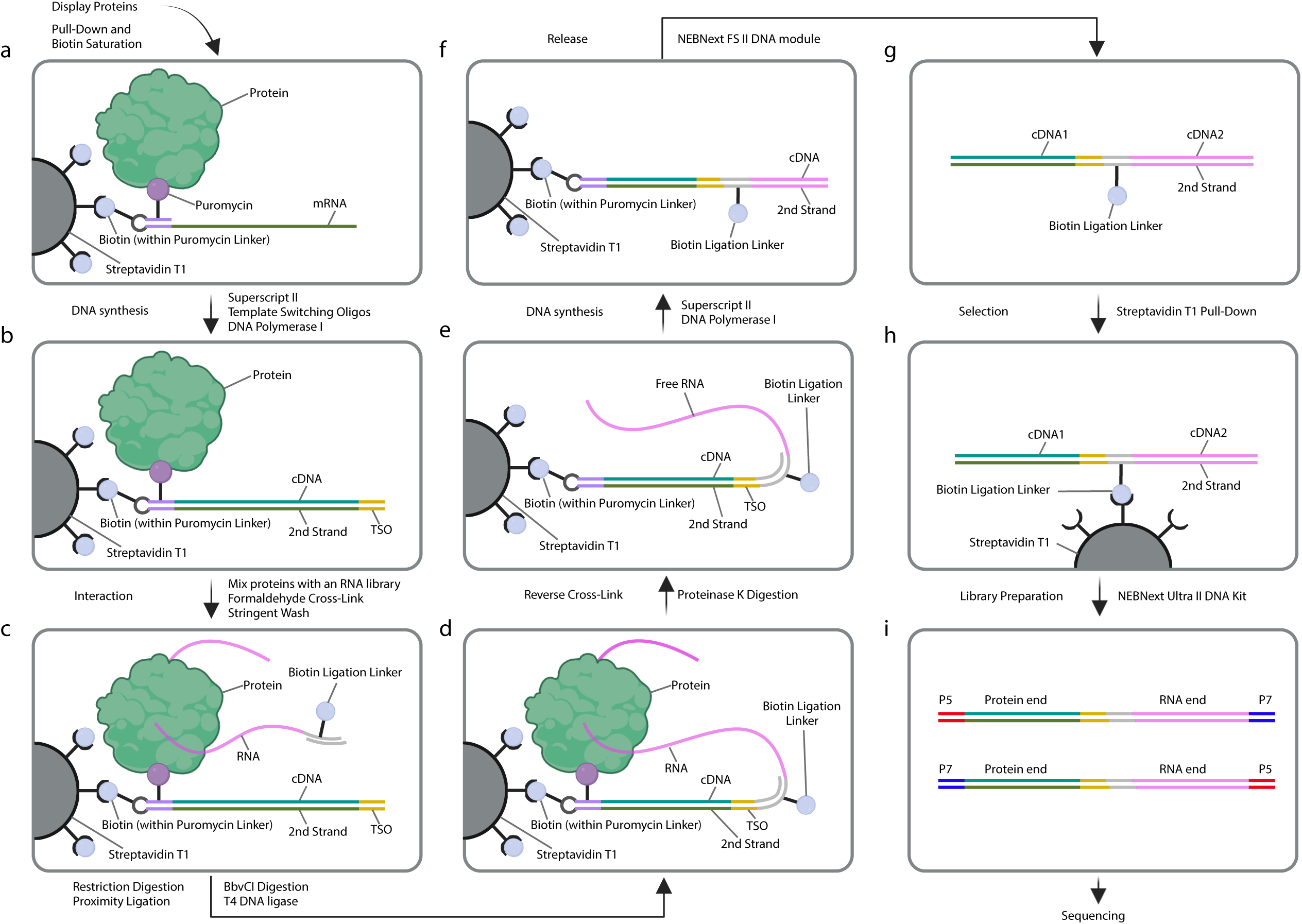
The RELIS procedure. (a) The library of mRNA-labeled proteins is immobilized. (b) the mRNA labels are converted to cDNA. (c) An RNA library is incubated with the protein library to allow for interactions. (d) The RNA (pink) is ligated with the protein’s cDNA label (green) via a linker sequence (gray), creating a chimeric sequence in the form of RNA-linker-cDNA, which is converted to double-stranded DNA for paired-end sequencing (e-i).

The bioinformatics module resolves the RNA-protein pairs from the chimeric sequences (Figure 1c). This module performs three main functions. First, it identifies the gene pair mapped with the two reads of a read pair. Second, it determines the protein-end and the RNA-end. Our experimental design ensures that any sequencing read from the 5’ end is either the sense strand of the RNA or the antisense strand of the cDNA (Figure S2a). Leveraging this fact, the bioinformatics module assigns the read mapped to the sense strand as the RNA-end and the read mapped to the antisense strand as the protein-end. Read pairs where both reads are mapped to the sense or the antisense strand are excluded from downstream analysis. The retained read pairs are termed “chimeric read pairs.”

The third function of the bioinformatics module is to identify associating RNA-protein pairs using statistical tests. The input for these tests is the set of “chimeric read pairs.” The null hypothesis for each gene pair is that the presence of reads originating from one gene is independent of the presence of reads originating from the other gene. A Chi-square test is performed for each gene pair (Figure S2b), and Bonferroni-Hochberg (BH) correction is applied to account for multiple hypothesis testing ^49^. An RNA-protein pair is identified when the BH-corrected p-value is less than 0.05 and the number of read pairs mapped to this gene pair is at least four times the expected number of read pairs (number of read pairs > 4X, where X is the expected number of read pairs). We have implemented these data processing and statistical analyses into an open-source software package called PRIMseqTools (Figure S2c), which is available at <u>PRIMseqTools GitHub</u> <u>repository</u>.

### A human RNA-protein association network (HuRPA)

To derive an RNA-protein association network from human cells, we generated PRIM-seq libraries from human embryonic kidney cells (HEK293T) and human lymphoblast cells (K562) (Table S1). Comparison of biological replicate libraries within HEK293T and K562 revealed significant overlaps between replicate libraries (odds ratio > 2241.3, p-value < 1e-323, Chi-square test, Figure S3). Furthermore, the overlaps increased as the threshold for calling RNA-protein associations was raised (Figure S3), indicating good reproducibility of these replicate libraries.

We merged the sequencing data from all the PRIM-seq libraries into a single PRIM-seq dataset. Applying PRIMseqTools to this dataset revealed 365,094 RNA-protein associations (BH-corrected p-value < 0.05 and number of read pairs > 4X), involving 11,311 proteins and 7,248 RNAs (Figure 3a, Figure S4a,b). We call this network the <u>Hu</u>man <u>R</u>NA-<u>P</u>rotein <u>A</u>ssociation Network (HuRPA), and refer to its involved proteins and RNAs as HuRPA proteins and HuRPA RNAs. We developed a web interface to query, visualize, and download HuRPA (https://genemo.ucsd.edu/prim).

**Figure 3.**
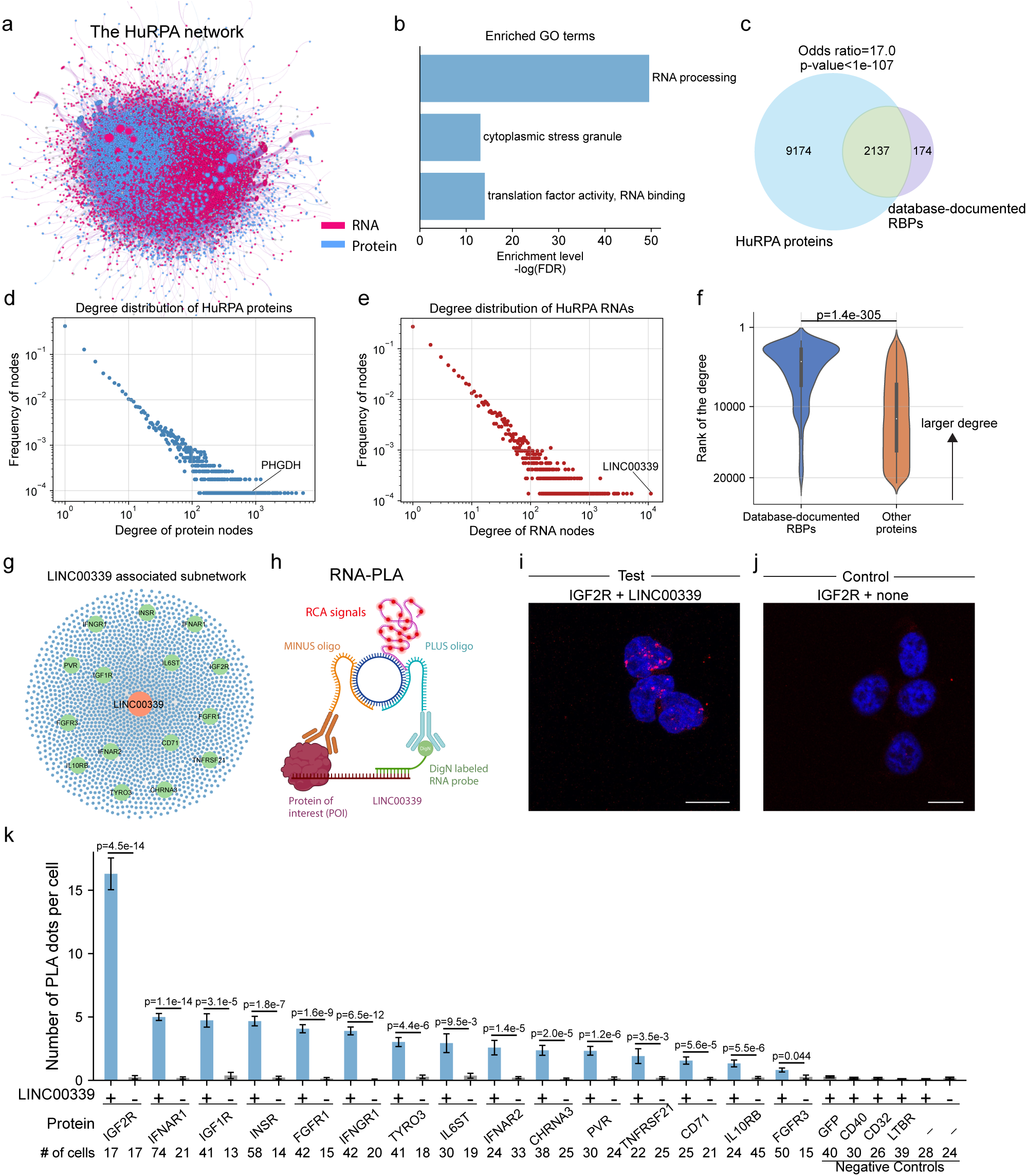
HuRPA network. (a) The entire HuRPA network with proteins (blue) and RNAs (red) as nodes, RNA-protein associations as edges, and the node size representing this node’s number of edges (degree). (b) The most enriched GO terms in HuRPA proteins. X axis: -log_e(FDR). (c) Overlap between HuRPA proteins and database-documented RBPs. (d-e) degree distribution of HuRPA proteins (d) and HuRPA RNAs (e) with the degree on the x axis and the frequency of the nodes with the corresponding degree on the y axis. (f) Rank distribution of the degrees of database-documented RBPs (blue) and the other proteins (orange) in HuRPA. The protein with most associated RNAs (highest degree) is ranked on top (rank = 1). (g) The RNA hub, LINC00339 and its associated proteins (green nodes). The 15 to-be-tested proteins are highlighted. (h) Illustration of the RNA-Proximity ligation (RNA-PLA) assay. RCA: rolling circle amplification. (i-j) representative microscopic images of the RNA-PLA assay on the IGF2R protein and LINC00339 RNA (i) and antibody-only control (j). Scale bar = 20 μm. (k) Quantification of 15 tests (LINC00339 +, protein = IGF2R, …, FGFR3), antibody-only controls (LINC00339 -, protein = IGF2R, …, FGFR3), other negative controls including LINC00339(RNA)-GFP(protein), LINC00339(RNA)-CD40(protein), (RNA)-CD32(protein), (RNA)-LTBR(protein), RNA probe-only control (LINC0039 +, Protein -); and no-probe-no-antibody control (LINC0039 -, Protein -). # of cells: the number of cells used for quantification. Y axis: the average number of RNA-PLA foci per cell. Error bar: SEM.

We tested whether the HuRPA proteins are enriched with any Gene Ontology (GO) annotations^50^. “RNA processing” (GO:0006396), “cytoplasmic stress granule” (GO:0010494), and “Translation factor activity, RNA binding” (GO:0008135) emerged as the most enriched BP, CC, and MF terms, respectively (FDR = 2.99e-22, 2.09e-6, and 7.99e-7, Fisher’s exact test) (Figure 3b, Figure S5). The “RNA processing” associated HuRPA proteins included RNA splicing factors, RNA metabolism proteins, RNA processing factors, RNA methyltransferases, ribosomal proteins, and RNA helicases (Figure S5c). “Cytoplasmic stress granules” are condensates of proteins and RNAs (Figure S5d,e) (Protter and Parker 2016). These data highlight RNA-protein association as the most prominent characteristic of HuRPA among all functional annotations.

We compared HuRPA proteins with database-documented RBPs. We compiled RBPs from six databases: RBP2GO ^51^, RBPDB ^52^, ATtRACT ^53^, hRBPome ^54^, RBPbase ^55^, and starBase ^56^, resulting in 2,311 database-documented RBPs (Figure S4c). Most of these (2,137, 92.5%) are HuRPA proteins, representing 18.9% of the 11,311 HuRPA proteins and suggesting a significant overlap (odds ratio = 17.0, p-value = 3.e-108, Chi-square test, Figure 3c). Thus, HuRPA recovers most database-documented RBPs.

We explored whether any HuRPA proteins outside the database-documented RBPs (Undatabased HuRPA proteins) have been detected as candidate RBPs by recent technologies. Among the 9,174 Undatabased HuRPA proteins, 764 were captured by peptide cross-linking and affinity purification (pCLAP) (Mullari et al. 2017) (odds ratio = 8.1, p-value = 3.1e-89, Chi-square test) and 129 by RBDmap (Castello et al. 2016) (odds ratio = 18.2, p-value = 1.1e-66, Chi-square test), revealing enrichments of the Undatabased HuRPA proteins in the candidate RBPs detected by recent technology (Figure S4d, Table S2). These data corroborate the idea that PRIM-seq can detect previously uncharacterized RBPs.

We compared database-documented RNA-protein association (RPA) pairs with HuRPA’s RNA-protein association pairs (HuRPA RPAs). Using the RPAs documented in the RNAInter database^57^ as the reference set, HuRPA RPAs exhibited significantly higher precision and recall than randomly sampled RPAs (Figure S6a). Furthermore, as we increased the read count threshold for calling RPA from the PRIM-seq data, the resulting subnetworks of HuRPA exhibited larger precision, i.e. a larger fraction of the subnetwork being RNAInter RPAs (Figure S6a). Changing the reference set to RPAs detected by iCLIP (iCLIP RPAs) or HITS-CLIP (HITS-CLIP RPAs) (Table S2) revealed similar outcomes, where HuRPA RPAs exhibited significantly higher precision and recall than randomly sampled RPAs (Figure S6b,c). These data suggest a consistency between HuRPA RPAs and previously identified RPAs.

HuRPA exhibits a scale-free topology ^58^, where the number of proteins is negatively correlated with the number of their associated RNAs (Figure 3d), and conversely, the number of RNAs is inversely correlated with the number of their associated proteins (Figure 3e). We asked whether a HuRPA protein with more associated RNAs, i.e. a higher degree is more likely to be detected by other methods. Database-documented RBPs exhibited higher degrees in HuRPA than the other proteins (p-value = 1.4e-305, t-test, two-sided, Figure 3f). Furthermore, among the Undatabased HuRPA proteins, those detected by either pCLAP or RBDmap exhibited higher degrees in HuRPA than the remaining Undatabased HuRPA proteins (p-value = 4.8xe-10 for pCLAP, p-value = 1.1e-59 for RBDmap, t-test, two-sided, Figure S4e). Thus, the more associated RNAs a protein has in HuRPA, the more likely it is detected by another technology.

Moreover, we obtained highly connected subnetworks of HuRPA by removing the HuRPA proteins with small degrees (i.e., small numbers of associated RNAs). As the degree threshold for removing proteins increased, the subnetworks exhibited higher precision and recall compared to RNAInter RPAs as the reference set (Figure S6d). Changing the reference set from RNAInter RPAs to iCLIP RPAs and HITS-CLIP RPAs led to similar results (Figure S6e,f). Thus, the HuRPA proteins with more associated RNAs are more likely detected by other methods.

### Enrichment of RNA binding domains and RBD-binding motifs in PRIM-seq reads

When we designed PRIM-seq, we did not anticipate a strong correlation between PRIM-seq reads and the RNA-binding domains (RBDs) within proteins. Nevertheless, we compared PRIM-seq’s protein-end reads with the protein sequences. HuRPA includes 1,831 RBD-containing proteins, with a total of 3,702 RBDs. These RBDs represent 2.1% of the total length of the mature mRNAs of these RBD-containing proteins. Remarkably, 13.9% of the protein-end reads were mapped to these RBDs, indicating a significant enrichment of PRIM-seq protein-end reads originating from RBDs (p-value = 4.1e-272, binomial test, one-sided, Figure S7a). For example, the Heterogeneous nuclear ribonucleoprotein R (HNRNPR) has four RBDs, which align with regions of abundant protein-end reads (Figure S7b).

HuRPA proteins contain seven classes of RBDs: RNA-Recognition Motifs (RRMs), PseudoUridine synthase and Archaeosine transglycosylase (PUA) domains, Like Sm (LSm) domains, K Homology (KH) domains, Cold-shock Domains (CSDs), DEAD domains, and ribosomal protein-S1-like (S1) domains ^52^ (Table S3). Each RBD class showed enrichment in PRIM-seq’s protein-end reads (largest p-value = 8.0e-18, binomial test, one-sided, Figure S7c,d, Table S3).

RNA sequences bound by an RBD often exhibit conserved sequence patterns, known as RBD-binding motifs ^59^. We focused on RRMs for subsequent analysis, as RRM is the largest RBD class and the most enriched with protein-end reads (p-value = 8.6e-319, binomial test, one-sided). We retrieved the RNA-end reads that paired with the RRM-mapped protein-end reads. *De novo* motif finding revealed 13 RNA sequence motifs (motif length = 10, BH-corrected p-value < 0.05), three of which matched previously reported RNA-recognition sequences for RRM ^53^ (Figure S7e). These data suggest that PRIM-seq’s protein-end and RNA-end reads are enriched with RBDs and RBD-binding motifs, respectively.

### Experimental validation of RNA-protein associations

We set out to validate previously uncharacterized RNA-protein associations (RPAs) in HuRPA, beginning with the long intergenic noncoding RNA (lincRNA) LINC00339. LINC00339 is the HuRPA RNA with the largest number of associated proteins, making it the largest RNA hub (Figure 3e,g). Notably, LINC00339 has no previously reported interacting proteins. It is expressed in most human tissues and is associated with tumorigenesis and progression of various cancers ^6061^. Given LINC00339’s wide distribution across most subcellular compartments, including the cell membrane ^62^, we selected 15 LINC00339-associated proteins from HuRPA for validation, representing diverse subcellular compartments (green nodes in Figure 3g, Table S4).

We used the RNA-proximity ligation assay (RNA-PLA) for validation ^63^. RNA-PLA detects specific RNA-protein interactions by using an RNA probe targeting the specific RNA and an antibody recognizing the associated protein (Figure 3h). As a sanity check, we reproduced the contrast of RNA-PLA signals between the U1 snRNA-Smith protein complex and the Smith antibody-only control ^64^ in HEK293T cells (Figure S8).

We applied RNA-PLA to the 15 selected RNA-protein pairs: LINC0039 with IGF1R, IGF2R, CHRNA3, INSR, TYRO3, CD71, IL6ST, IL10RB, FGFR1, FGFR3, IFNAR1, IFNAR2, IFNGR1, PVR, and TNFRSF21 (Figure 3i, Figure S9a, Table S4). We included four sets of controls: antibody-only controls (RNA probe not added); four RNA-protein pairs not included in HuRPA (LINC0039-CD40, LINC0039-CD32, LINC0039-LTBR, and LINC0039-green fluorescent protein (GFP)); RNA probe-only control (antibody not added); and no-probe-no-antibody control (Figure 3j, Figure S9b,c). The controls consistently exhibited few RNA-PLA foci (Figure 3j, Figure S9b,c), while each tested RNA-protein pair showed significantly more RNA-PLA foci per cell than the controls (smallest p-value = 1.1e-14 for IFNAR1, largest p-value = 0.044 for FGFR3, t-test, two-sided, Figure 3k, Figure S9). For example, the LINC0039-IGF2R pair exhibited an average of 16.3 RNA-PLA foci per cell, which is 69-fold higher than the antibody-only control (p-value = 4.5e-14, t-test, two-sided) (Figure 3i-k). These data suggest that previously uncharacterized RNA-protein associations in HuRPA can be validated using alternative methods.

### Phosphoglycerate dehydrogenase (PHGDH) as an RNA-associating protein

We proceeded to test an Undatabased HuRPA protein that has not been documented in any RBP database ^5256545351^. We chose Phosphoglycerate dehydrogenase (PHGDH) for validation, as it is an Undatabased HuRPA protein and a hub protein associated with a large number of RNAs (Figure 3d, Figure 4a, Figure S10a). Additionally, PHGDH appears to have conflicting roles compared to its known function as an enzyme in the serine synthesis pathway ^65^ in some neural systems ^66^, suggesting the possibility of an uncharacterized function.

**Figure 4.**
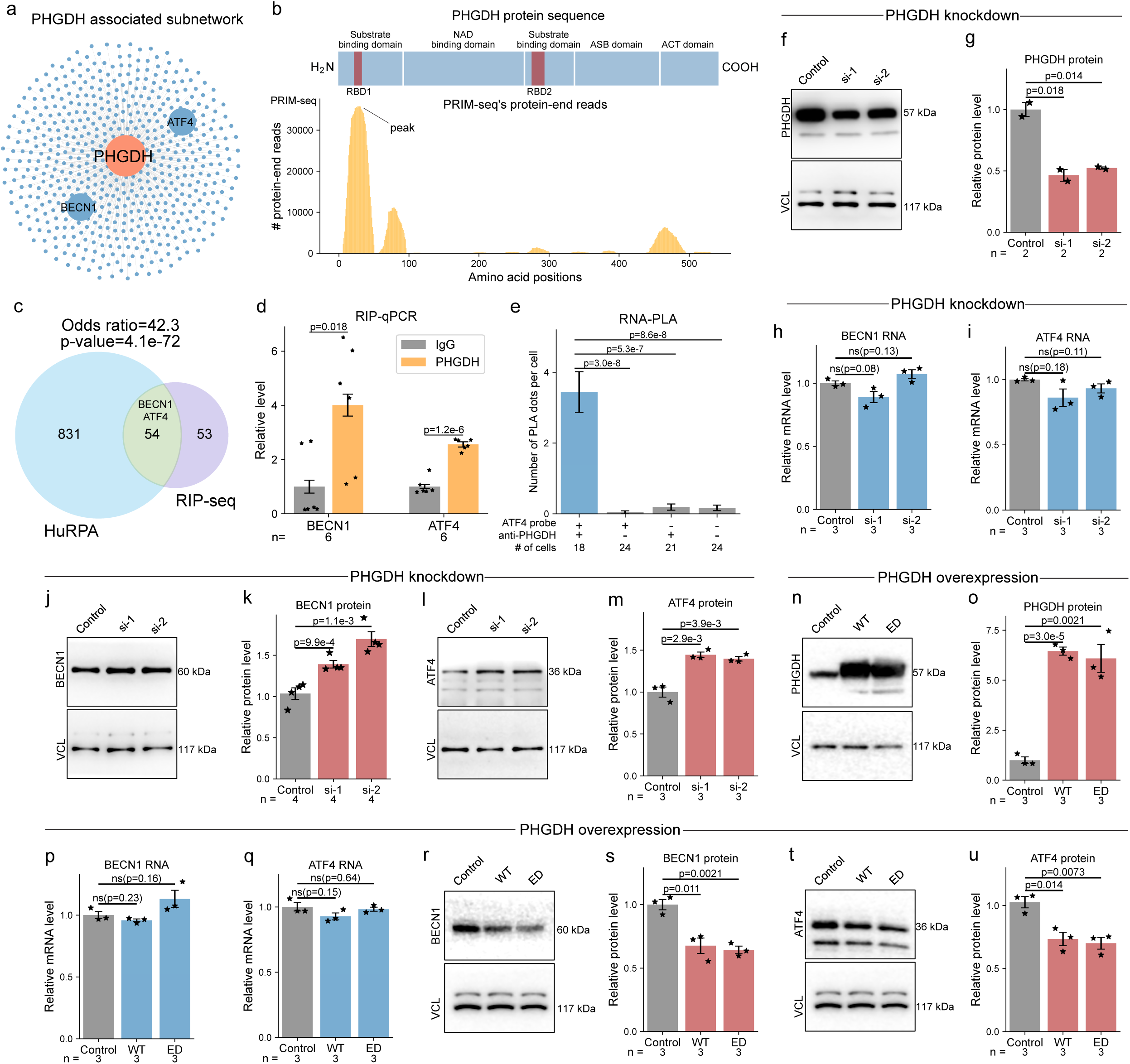
PHGDH as an RNA-associating protein. All p-values are derived from two-sided t tests and error bars represent SEM unless otherwise specified. The number of replicates is denoted as “n=” in the bottom row. (a) The protein hub, PHGDH, and its 885 associated RNAs (blue nodes). (b) Upper panel: the database-documented protein domains (blue blocks) and the two RBDmap-identified candidate RNA-binding domains (RBD1, RBD2) in PHGDH. Lower panel: PRIM-seq’s protein-end reads (yellow) aligned to PHGDH’s peptide sequence, with the peak co-localizing with RBD1. (c) Overlap of the HuRPA and RIP-seq identified PHGDH-associated RNAs. P-value is derived from Fisher’s exact test. (d) RIP purified BECN1 and ATF4 RNA levels using PHGDH antibody (yellow) and IgG (gray). n=6. (e) RNA-PLA analysis with and without the ATF4 RNA probe (ATF4 probe = +, -), with and without the PHGDH antibody (anti-PHGDH = +, -). The average number of PLA foci per cell (y axis) is derived from the number of cells given in the bottom row (# of cells). (f-m) PHGDH knockdown with PHGDH targeting siRNAs (si-1, si-2) and a scramble siRNA (Control). (f) si-1, si-2 reduced PHGDH protein level (f-g), without changing the mRNA levels of BECN1 (h) and ATF4 (i). ns: non-significant. Western blotting (j, l) and quantification (k,m) of BECN1 and ATF4 with Vinculin (VCL) as the loading control. PHGDH knockdowns si-1 and si-2 induced protein levels of BECN1 (j-k) and ATF4 (l-m). (n-u) Overexpression of wild-type (WT) and enzymatically-dead (ED) PHGDH and without overexpression (Control). Western blots (n) and quantification (o) showing the overexpression of WT and ED PHGDH, without affecting the mRNA levels of BECN1 (p) and ATF4 (q), which suppress the protein levels of BECN1 (r-s) and ATF4 (t-u).

Consistent with PRIM-seq data, a proteome-wide RNA-binding domain (RBD) screening by peptide-mass spectrometry (RBDmap) reported two candidate RBDs within the PHGDH protein (Castello et al (2016) (RBD1 and RBD2, Figure 4b). Importantly, more than 60% of PRIM-seq’s protein-end reads mapped to PHGDH formed a peak (36,210 in-peak reads / 59,986 total reads), pinpointing RBD1 (AA 22-33) (p-value = 2.6e-214, binomial test, one-sided, Figure 4b). This data highlights the consistency between PRIM-seq, a DNA-sequencing-based technology, and RBDmap, a peptide-mass spectrometry-based technology.

To further test if PHGDH is an RNA-associating protein, we performed RNA immunoprecipitation followed by sequencing (RIP-seq) on PHGDH with IgG as the control in HEK293T cells in two biological replicates (Table S5). The RIP-seq data revealed 107 PHGDH-associated RNAs, 54 of which overlapped with PHGDH-associated RNAs in HuRPA, showing a strong enrichment (odds ratio = 42.3, p-value = 4.1e-72, Fisher’s exact test, Figure 4c, Figure S10b). Moreover, as we increased the threshold for calling PHGDH-associated RNAs from RIP-seq, the degree of overlap with PHGDH-associated RNAs in HuRPA, indicated by the odds ratio, also increased (Figure S10c). Conversely, PHGDH-associated RNAs with higher read counts in HuRPA were more often detected by RIP-seq (p-value = 0.034, t-test, two-sided, Figure S10d). These RIP-seq data corroborate PHGDH’s RNA association ability and validate a subset of PHGDH-associated RNAs identified by PRIM-seq.

The mRNAs of Beclin-1 (BECN1) and Activating Transcription Factor 4 (ATF4) are among the PHGDH-associated RNAs in both HuRPA and the RIP-seq identified targets (Figure 4c, Figure S10b). Considering BECN1 and ATF4’s roles in regulating autophagy and apoptosis ^67,686970^, we specifically tested the association of these two mRNAs with PHGDH. RIP followed by quantitative PCR (RIP-qPCR) confirmed the co-precipitation of BECN1 mRNA (p-value = 0.018, t-test, two-sided) and ATF4 mRNA with PHGDH (p-value = 1.2e-6 for ATF4, t-test, two-sided, Figure 4d). Additionally, we used RNA-PLA to test the ATF4 mRNA-PHGDH association. The ATF4 mRNA-PHGDH pair exhibited 18-85 fold more PLA foci compared to the three controls lacking the RNA probe (p-value = 5.3e-7), the antibody (p-value = 3.0e-8), or both the RNA probe and the antibody (p-value = 8.6e-8, t-test, two-sided, Figure 4e, Figure S10e-h). This data further supports the association of PHGDH protein with BECN1 and ATF4 mRNAs.

### PHGDH regulates the protein levels of its associated mRNAs

We tested whether PHGDH modulates either the mRNA or protein levels of BECN1 and ATF4. Two PHGDH-targeting siRNAs (si-1, si-2) reduced PHGDH levels by approximately 50% compared to a scrambled siRNA (Control) in HEK293T cells (p-value = 0.018 for si-1, p-value = 0.014 for si-2, t-test, two-sided, Figure 4f,g), confirming effective knockdown of PHGDH. Neither siRNA altered the mRNA levels of BECN1 or ATF4 (p-values > 0.05 for both mRNAs, t-test, two-sided, Figure4h,i). However, both siRNAs increased the protein levels of both BECN1 and ATF4 (p-value < 0.0011 for BECN1, p-value < 0.0039 for ATF4, t-test, two-sided, Figure 4j-m).

The induction of BECN1 protein is expected to enhance autophagy and impair cell proliferation ^67,68^. Consistent with these expectations, immunostaining analysis revealed increases in autophagosome formation in both PHGDH knockdowns compared to the scramble control in HEK293T cells (p-value = 4.1e-4 for si-1, p-value = 4.7e-4 for si-2, t-test, two-sided, Figure S11a,b). Furthermore, Bromodeoxyuridine (BrdU) incorporation decreased in both knockdowns, suggesting reduced cell proliferation (p-value = 3.8e-7 for si-1, p-value = 1.2e-4 for si-2, t-test, two-sided, Figure S11c,d). Additionally, in mouse neural stem cells (mNSCs), two PHGDH-targeting siRNAs increased autophagosome levels (p-value = 3.8e-3 for si-1, p-value = 1.4e-3 for si-2, t-test, two-sided, Figure S12b,c) and reduced cell proliferation compared to the scramble control (p-value = 1.6e-2 for si-2, t-test, two-sided, Figure S12d,e), suggesting that these functional consequences are shared across several cell types and between humans and mice.

The induction of ATF4 protein is expected to promote apoptosis and neurite outgrowth ^6970^. Immunostaining analysis revealed increased activated Caspase-3 (aCaspase3), an apoptosis marker, in both knockdowns compared to the scramble control in HEK293T cells (p-value = 2.1e-6 for si-1, p-value = 9.7e-4 for si-2, t-test, two-sided, Figure S11e,f) and also in mESCs (p-value = 0.031 for si-1, p-value = 0.0099 for si-2, t-test, two-sided, Figure S12f,g). Furthermore, Sholl analysis of the morphology of mNSCs revealed longer dendrites and more dendritic crossings in the knockdowns compared to the scramble control (p-value = 0.018 for si-1, p-value = 0.0071 for si-2, Kolmogorov–Smirnov test, two-sided, Figure S12h-j). Thus, reducing PHGDH promotes apoptosis in both cell types and neurite outgrowth in mNSCs.

To determine whether these protein level changes of BECN1 and ATF4 can be attributed to PHGDH’s enzymatic function, we overexpressed wild-type (WT) PHGDH and an enzymatically-dead (ED) variant of PHGDH in HEK293T cells (Figure 4n,o). The ED variant abolishes PHGDH’s enzymatic function with an R236Q mutation on the active site ^71^. Overexpression of both WT and ED PHGDH reduced the protein levels of BECN1 and ATF4 (p-value < 0.011 for BECN1, p-value < 0.014 for ATF4, t-test, two-sided, Figure 4r-u) without discernible changes in their mRNA levels (p-values > 0.05 for both BECN1 and ATF4, t-test, two-sided, Figure 4p,q). These overexpression data corroborate PHGDH’s suppressive roles in BECN1 and ATF4 protein expression and suggest this role is independent of PHGDH’s enzymatic function. Taken together, these examples highlight PRIM-seq’s ability to unveil uncharacterized RNA-protein associations.

## Discussion

The advent of PRIM-seq (Protein-RNA Interaction Mapping by sequencing) represents a transformative step in the study of RNA-protein associations. By allowing for the high-throughput identification of RNA-associating proteins and their associated RNAs, PRIM-seq addresses the limitations of existing methodologies that often require specific reagents and are constrained by a one-to-many mapping approach. Our study demonstrates the efficacy of PRIM-seq through the construction of the human RNA-protein association network (HuRPA), which encompasses 7,248 RNAs, 11,311 proteins, and 365,094 RNA-protein pairs, significantly expanding the known landscape of these associations.

A compelling aspect of PRIM-seq is its ability to perform genome-wide mapping without prior knowledge of specific RNA or protein targets. This capability is particularly valuable given the sheer number of potential RNA-protein pairs within the human genome, estimated to involve approximately 60,000 coding and 30,000 noncoding genes. By converting RNA-protein associations into unique chimeric DNA sequences for high-throughput sequencing, PRIM-seq efficiently deciphers these complex networks at an unprecedented scale.

The HuRPA network revealed by PRIM-seq exhibits a scale-free topology, characterized by a few hub proteins interacting with many RNAs and a larger number of proteins with fewer RNA partners. This finding is consistent with known architecture of other biological networks. The significant overlap of HuRPA proteins with database-documented RBPs supports the reliability of our approach. Interestingly, database-documented RBPs are even more enriched in the hub proteins of the HuRPA network, suggesting the proteins with more RNA partners are more likely revealed by multiple experimental techniques.

Our experimental validations further underscore the robustness and utility of PRIM-seq. We identified LINC00339 as a non-coding RNA with extensive protein associations and validated its interactions using RNA-proximity ligation assay (RNA-PLA). This non-coding RNA’s association with diverse proteins, many of which are implicated in various cellular compartments and processes, suggests a broad regulatory role that warrants further investigation. Additionally, the identification of PHGDH as an RNA-associating protein that modulates the protein levels of BECN1 and ATF4 illustrates the functional impact of these interactions. The suppression of BECN1 and ATF4 protein expression by PHGDH reveals a novel regulatory mechanism influencing autophagy, apoptosis, and cell proliferation.

Notably, while PRIM-seq is designed to capture RNA-protein complexes rather than direct binding events, the enrichment of RNA-binding domains (RBDs) in the protein-end reads and RNA-recognition motifs in the RNA-end reads suggests that many of the detected interactions may indeed involve direct binding. In conclusion, PRIM-seq offers a powerful and scalable solution for the comprehensive mapping of RNA-protein associations. The extensive HuRPA network and the newly validated associations demonstrate the potential of PRIM-seq to advance our understanding of RNA biology and its regulatory mechanisms.

### Online Methods Cell culture

HEK293T (ATCC, CRL3216) cells were cultured in Dulbecco’s modified Eagle medium (DMEM; Gibco™, 11960044) supplemented with 10% fetal bovine serum (FBS) (Gemini, 100-500), 2 mM Glutamax (Gibco™, 35050061), and 5,000 U/ml penicillin/streptomycin (Gibco™, 15070063), at 37°C with 5 % CO2. K562 (ATCC, CCL-243) cells were cultured in Iscove’s Modified Dulbecco’s Medium (IMDM) (ATCC, 30-2005) supplement with 10% fetal bovine serum (FBS) (Gemini, 100-500), at 37°C with 5 % CO2.

Mouse neural stem cells (mNSCs) were collected from the fetal forebrains of a mouse (C57BL/6J, JAX Lab, Strain #:000664, RRID:IMSR_JAX:000664) embryo at Day 16. The brain was crosscut followed by a trypsin digestion (Gibco™, 25200056) and a cell strainer (70 µm Nylon) filtration (PlutiSelect, 43-50070-51) to obtain single cells. The mNSC basal medium including DMEM/F12 (Gibco™, 11320033) with 1% L-glutamine (Corning®, 10-090-CV), 2% B27 without VA (Gibco™, 12587010), and 1% penicillin/streptomycin (5000 U/ml, Gibco™, 15070063) was used to culture mNSCs in the presence of 20 pg/µL epidermal growth factor (EGF; PeproTech, 100-15) and basic fibroblast growth factor-2 (FGF-2; PeproTech, 100-18B-B). Cell densities were checked every day. If the cell density was high, a subculture was performed; if not, the medium was changed by replacing half of the overnight medium to a fresh medium. During subculture, cells were collected from a 6 cm dish into a 15 mL tube. The cell culture was centrifuged at 1.5k rpm for 2 minutes. The supernatant was discarded, and cells were resuspended using 140 μL of NSC basal medium with growth factors. Approximately 110 μL of the cell suspension was discarded, and the rest of the cell suspension was dispersed in 4 mL of NSC basal medium with growth factors into a new 6 cm dish.

## PRIM-seq

### Preparation of the linker

The linker is a critical reagent. It is designed for efficient ligations with the RNA on one end and with the protein’s cDNA label on the other end, to create a RNA-linker-cDNA structure. The linker is composed of a top strand oligo (5’-/Phos/TGACCAATGGCGCCGGGCCTTTCTTTATGTTTTTGGCGTCTTGG-3’, IDT) and a biotinylated bottom strand oligo (5’-/Phos/TGACCAAGACGCCAAAAACA/iBiodT/AAAGAAAGGCCCGGCGCCATTGG-3’, IDT). The two strands were separately dissolved in 200 μM with UltraPure™ DNase/RNase-Free distilled water (Thermo Scientific, 10977023). The bottom strand was adenylated with the 5’ DNA Adenylation Kit (NEB, E2610S) to create an “App-bottom-strand” (5’-/App/TGACCAAGACGCCAAAAACA/iBiodT/AAAGAAAGGCCCGGCGCCATTGG-3’) and purified with Zymo ssRNA/DNA Clean & Concentrator Kit (Zymo Research, D7010).

### SMART-Display

SMART-display was carried out as previously described to create a library of mRNA-labeled proteins ^48^.

### REILIS (<u>Re</u>verse transcription, Incubation, <u>L</u>igation, and <u>S</u>equencing)

REILIS includes 3 steps. **Overview of Step 1:** The first step is the incubation of the SMART-display protein library with an RNA library. In this step, displayed protein libraries are immobilized on streptavidin T1 beads, and their mRNA labels are reverse transcribed to double-stranded cDNA with a template-switching oligo (TSO) that contains a BbvCI restriction site. The TSO was digested with BbvCI. Next, RNA is extracted from input cells, ligated with the linker, and incubated with the protein library, allowing for RNA-protein interactions. The linker has a sticky end complementary to the BbvCI site on the TSO, poised for efficient ligation.

Protein immobilization: 100 uL of Dynabeads MyOne Streptavidin T1 Beads (Invitrogen™, 65602) were prepared according to the manufacturer’s recommendations. The SMART-display proteins were incubated with the suspended beads for 1 hour with rotation at room temperature. Next, 50 uM free biotin (Invitrogen, B20656) was incubated with the streptavidin beads for 10 mins to block the remaining binding sites. The beads were washed three times with 1x PBS pH 7.4 (Gibco™, 70011044) with 0.1% Triton X-100 (Sigma-Aldrich, T8787-50ML).

Conversion of protein’s mRNA label to cDNA: 50 μL of first-strand synthesis mix was created with 500 U of SuperScript II Reverse Transcriptase (Thermo Scientific, 18064014), 1x SuperScript II FS Buffer (Thermo Scientific, 18064014), 5 mM DTT (Thermo Fisher Scientific, P2325), 1 uM dNTP mix (NEB, N0447S), 1 M Betaine (Sigma-Aldrich, 61962), 6 mM MgCl2 (Thermo Scientific, R0971), 500 pmol of End Capture TSO (5’-/dSp/AGT AAA GGA GAC CTC AGC TTC ACT GGA rGrGrG-3’, IDT), and 40 U of SUPERase• In™ RNase Inhibitor (Invitrogen™, AM2694). The mix was incubated with the protein-bound beads at 42°C for 50 minutes with agitation, and then cycled 10 times at 50°C for 2 minutes followed by 42°C for 2 minutes. The beads were washed twice for 5 minutes with 500 μL 1x PBS pH 7.4 (Gibco™, 70011044) with 0.1% Triton™ X-100 (Sigma-Aldrich, T8787-50ML). To synthesize the second-strand cDNA, 100 μL of second-strand synthesis mix was created with 20 U DNA Polymerase I (NEB, M0209S), 1x NEBuffer 2 (NEB, M0209S), 2.4 mM DTT (Thermo Fisher Scientific, P2325), and 0.25 mM dNTP mix (NEB, N0447S). The mix was incubated with the protein-bound beads at 37°C for 30 minutes with agitation. The beads were washed twice for 5 minutes with 500 μL 1x PBS pH 7.4 (Gibco™, 70011044) with 0.1% Triton™ X-100 (Sigma-Aldrich, T8787-50ML).

Ligation of RNA and linker: The total RNA was extracted from approximately 10 million input cells with TRIzol reagent (Invitrogen, 15596026) according to the manufacturer’s recommendations. The purified RNA was fragmented with the NEBNext® Magnesium RNA Fragmentation Module (NEB, E6150S) for 2 minutes at 94 °C and purified with RNeasy Mini Columns (Qiagen, 74104). 200 pmols of RNA were end-repaired in a 200 μL reaction solution containing 100 U Quick CIP (NEB, M0525S) and 1x rCutSmart buffer (B6004S) at 37 °C for 1 hour and then purified with RNeasy Mini Columns (Qiagen, 74104). The end-repaired RNA was ligated with the adenylated bottom strand of the linker (App-bottom-strand) in 200 μL reaction mix, containing 400 pmols of App-bottom-strand, 200 pmols of RNA, 4,000 U T4 RNA Ligase 2, truncated KQ (NEB, M0373S), 1x T4 RNA Ligase buffer (NEB, M0373S), and 15% PEG 8000 (NEB, M0373S), at 16 °C overnight. The ligation product (RNA-bottom_strand) was purified with RNeasy Mini Columns (Qiagen, 74104). The top strand of the linker was annealed with the RNA-bottom_strand by mixing the top strand and RNA-bottom_strand at 1:1 molar ratio in 1x Annealing buffer [10x: 100 mM Tris-HCI Buffer, pH 7.5 (Invitrogen, 15567027), 500 mM NaCl (Thermo Scientific, AM9759), 10 mM EDTA pH 8.0 (Invitrogen™, AM9260G)], heating to 75°C for 5 minutes and cooling to 25°C at 0.1°C per second. This process creates a double-stranded linker with a sticky end that is complementary to the BbvCI restriction site on one side and with RNA ligated to its bottom strand on the other side (Figure S13).

Incubation of RNA and proteins: The protein-bound beads were suspended in 200 μL of RNA Binding Buffer (10 mM HEPES (Thermo Fisher Scientific, BP299100), 50 mM KCl (Invitrogen™, AM9640G), 4 mM MgCl2 (Thermo Scientific, R0971), 4 mM DTT (Thermo Fisher Scientific, P2325), 0.2 mM EDTA pH 8.0 (Invitrogen™, AM9260G), 7.6% glycerol (Invitrogen, 15514011)). This mix is added with 2 μg linker-ligated RNA and incubated at room temperature with rotation for 1 hour. Another 800 μL of Binding Buffer was added to bring the volume to 1 mL. The mix was rotated for an additional 10 minutes at room temperature.

**Overview of Step 2:** The second step ligates the protein’s cDNA label with the linker. This step crosslinks RNA-protein associations and applies stringent washes to remove non-complexed proteins or RNAs. The protein’s cDNA label is ligated with the linker via a sticky end ligation, creating a RNA-linker-cDNA structure. We note that the non-palindromic sticky end prevents self-ligation.

Crosslinking and washing: Crosslinking was performed at room temperature for 10 minutes at a final concentration of 1% formaldehyde (Thermo Fisher Scientific, 28906). The reaction was quenched with 125 mM glycine (Sigma-Aldrich, 67419-1ML-F) with rotation for 5 minutes. The beads were washed twice, each for 5 minutes with 500 μL Urea wash buffer [50 mM Tris-HCl pH 7.5 (Invitrogen™,15567027), 1% NP-40 (Thermo Scientific™, 85124), 0.1% SDS (Invitrogen™, AM9820), 2 mM EDTA pH 8.0 (Invitrogen™, AM9260G), 1 M NaCl (Thermo Fisher Scientific, AM9759), 4 M Urea (Sigma-Aldrich, U5378-1KG)], Low Salt wash buffer [0.1% SDS (Invitrogen™, AM9820), 0.1% Triton X-100 (Sigma-Aldrich, T8787-50ML), 2 mM EDTA pH 8.0 (Invitrogen™, AM9260G), 20 mM Tris-HCl pH 8 (Invitrogen™, 15568025), 150 mM NaCl (Thermo Fisher Scientific, AM9759)], and 1x PBS pH 7.4 (Gibco™, 70011044) with 0.1% Triton™ X-100 (Sigma-Aldrich, T8787-50ML).

Creating sticky end on the cDNA: The BbvCI site-containing TSO was treated with 10 U of BbvCI (NEB, R0601S) in 1x CutSmart Buffer at 500 μLs at 37°C for 1 hour with agitation. The beads were washed twice for 5 minutes each time with 500 μLs 1x PBS pH 7.4 (Gibco™, 70011044) with 0.1% Triton™ X-100 (Sigma-Aldrich, T8787-50ML).

Ligation of cDNA and linker: Proximity ligation was performed with 20,000 U of T4 DNA Ligase in 1 mL of 1x T4 DNA Ligase Buffer (NEB, M0202M), with constant rotation for 30 minutes at room temperature. The ligase was inactivated by heating to 65°C for 10 minutes. The beads were collected and washed twice for 5 minutes each time, with 500 μLs 1x PBS pH 7.4 (Gibco™, 70011044) with 0.1% Triton™ X-100 (Sigma-Aldrich, T8787-50ML).

**Overview of Step 3:** The third step constructs the sequencing library. This step converts the ligated RNA to double-stranded cDNA, denoted as cDNA2 to be differentiated from the protein’s cDNA label (cDNA1). The cDNA1-linker-cDNA2 chimeric sequence is released from beads, added with sequencing adapters, enriched by the biotin on the linker, amplified, and subjected to paired-end sequencing.

Protein digestion and reverse crosslinking: The streptavidin beads were suspended in 200 μL TAE buffer (Invitrogen™, AM9869) with 0.8 U of Proteinase K (NEB, P8107S) and incubated at 70°C for 30 minutes. The beads were washed twice for 5 minutes each time with 500 μL 1x PBS pH 7.4 (Gibco™, 70011044) with 0.1% Triton™ X-100 (Sigma-Aldrich, T8787-50ML).

Synthesis of cDNA2: 50 μL of first strand reaction mix was created with 500 U of SuperScript II Reverse Transcriptase, 1x SuperScript II FS Buffer, 5 mM DTT (Thermo Fisher Scientific, P2325), and 1 μM dNTP mix (NEB, N0447S), 1 M Betaine (Sigma-Aldrich, 61962), 6 mM MgCl2 (Thermo Scientific, R0971), and 40 U of SUPERase• In™ RNase Inhibitor (Invitrogen™, AM2694). The beads were incubated in this mix at 42°C for 50 minutes with agitation. The beads were washed twice for 5 minutes each time with 500 μL 1x PBS pH 7.4 (Gibco™, 70011044) and 0.1% Triton™ X-100 (Sigma-Aldrich, T8787-50ML). To synthesize the second strand cDNA, 100 μL of second strand mix was created with 20 U DNA Polymerase I (NEB, M0209S), 1 U RNase H (NEB, M0297S), 1x NEBuffer 2 (NEB, M0297S), 2.4 mM DTT (Thermo Fisher Scientific, P2325), and 0.25 mM dNTP mix (NEB, N0447S). The beads were incubated in this mix at 37°C for 30 minutes with agitation. The beads were washed twice for 5 minutes each time with 500 μL 1x PBS pH 7.4 (Gibco™, 70011044) with 0.1% Triton™ X-100 (Sigma-Aldrich, T8787-50ML).

Sequencing library construction: The cDNA-linker-cDNA2 structure was released from the beads by DNA fragmentation using the NEBNext® Ultra™ II FS DNA Module (NEB, E7810S) with twice the reaction volume and 10 minutes of fragmentation time. Sequencing adaptors were added using the NEBNext Ultra™ II DNA Library Kit (NEB, E7805S). The linker-containing DNA fragments were selected by incubating with 20 μL of Streptavidin T1 beads (Invitrogen™, 65602) at room temperature for 1h with agitation. The beads were washed 3 times, each time with a Low Salt wash buffer [0.1% SDS (Invitrogen™, AM9820), 0.1% Triton X-100 (Sigma-Aldrich, T8787-50ML), 2 mM EDTA pH 8.0 (Invitrogen™, AM9260G), 20 mM Tris-HCl pH 8.0 (Invitrogen™, 15568025), 150 mM NaCl (Thermo Fisher Scientific, AM9759)], 1x B&W buffer (5 mM Tris-HCl pH 7.5 (Invitrogen™, 15567027), 0.5 mM EDTA pH 8.0 (Invitrogen™, AM9260G), 1M NaCl (Thermo Fisher Scientific, AM9759)), and 1x PBS pH 7.4 (Gibco™, 70011044) with 0.1% Triton X-100 (Sigma-Aldrich, T8787-50ML). The beads were added to a PCR reaction mix consisting of 25 μL of 2x PCR Master Mix, 5 μL of Universal Primer, and 5 μL of Primer Mix-Index 1 from the NEBNext Ultra II Single Indexing Kit (NEB, E7335S). The PCR was conducted with an initial denaturation of 95°C for 2 minutes then cycled 15 times between a 98°C 10-second denaturation step and a 68°C 90-second annealing and extension step. PCR products were purified with 0.75x AMPure XP Beads (Beckman, A63880), eluted in 20 μL of UltraPure™ DNase/RNase-Free distilled water (Thermo Scientific, 10977023), and quantified with the Qubit dsDNA HS Assay Kit (Invitrogen™, Q32851). Each sequencing library was paired-end sequenced for 150 cycles on each end on an Illumina NovaSeq 6000 sequencer.

### RNA-PLA

Covalently coupling of DNA oligonucleotides to antibodies: The antibodies were conjugated with DNA oligonucleotides using a Duolink PLA Multicolor Probemaker Kit-Red (Sigma-Aldrich, DUO96010-1KT), adhering to the provided instructions in the manual.

RNA probes: The RNA probes targeting the RNA of interest are ultramer DNA oligonucleotides, synthesized by IDT DNA Technologies. Each RNA probe consists of three regions from 5′ to 3′, including a 40–50 nucleotide (nt) sequence complementary to (antisense to) the RNA of interest, 4 adenylates that serve as a linker, and a 3’ modification with Digoxin. An online FISH probe design resource was applied (e.g., http://prober.cshl.edu/) to identify region A for each target RNA. The sequences of the oligonucleotides used in this study are shown in Table S6.

Cell fixation and permeabilization: Approximately 5,000 HEK293T (ATCC, CRL3216) cells were subcultured in a Millicell EZ 8-well slide per well (Sigma-Aldrich, PEZGS0816). Once the cells reach to 70% - 90% confluence, culture medium was removed and the cells were fixed with 4% formaldehyde (v/v) (Thermo Scientific 043368.9M) in 1x PBS pH 7.4 (Gibco™, 70011044) on ice for 30 min. The cells were washed twice with 1x PBS pH 7.4 (Gibco™, 70011044) for 10 min each time. The cells were permeabilized with 200 µL of 0.1% Triton X-100 (Sigma-Aldrich, T8787-50ML) in 1x PBS pH 7.4 (Gibco™, 70011044) for 15 min at room temperature with rocking. Hybridization was blocked by incubation with 200 µL of blocking buffer (10 mM Tris-acetate pH 7.5 (BioWorld, 40125038), 10 mM magnesium acetate (Sigma-Aldrich, 63052-100ML), 50 mM potassium acetate (Sigma-Aldrich, 95843-100ML-F), 250 mM NaCl (Thermo Fisher Scientific, AM9759), 0.25 μg/μL bovine serum albumin [BSA] (Thermo Scientific, 23209), and 0.05% Tween 20 (Invitrogen™, AM9820)) in the presence of 20 μgs/mL sheared salmon sperm DNA (sssDNA) (Invitrogen™, AM9680) at 4°C for 1 h.

RNA probe hybridization: Ten nmols of RNA probes were diluted in 80 μL of UltraPure™ DNase/RNase-Free distilled water (Thermo Scientific, 10977023), denatured at 80 °C for 5 min, chilled on ice for 5 min, and resuspended in 80 μL of hybridization buffer (10% formamide (Thermo Scientific™, 17899), 2X SSC (Invitrogen™ 15557044), 0.2 mg/mL sheared salmon sperm DNA (Invitrogen™, AM9680), 5% dextran sulfate (Sigma-Aldrich, D8906-10G) and 2 mg/mL BSA (Thermo Scientific, 23209)), and incubated with the fixed and permeabilized cells at 37°C overnight to allow for hybridization. The cells were washed for 10 min twice with 2X SSC (Invitrogen™ 15557044) and twice with 1x PBS pH 7.4 (Gibco™, 70011044) at room temperature. Antibodies were blocked by incubation with 100 µL of Duolink Blocking Solution (Sigma-Aldrich, DUO92101-1KT) according to the manufacturer’s recommendations.

Incubation with antibodies: Two oligo-conjugated antibodies, including the antibody of protein of interest, which is conjugated with Oligo A from DUOLINK red kit (Sigma-Aldrich, DUO92008-100RXN), and anti-Digoxin, which is conjugated with Oligo B from DUOLINK red kit, (Sigma-Aldrich, DUO92008-100RXN), were mixed in Probemaker PLA Probe Diluent (Sigma-Aldrich, DUO82036) to a total volume of 200 µL and added to the fixed cells. The slides were incubated in a humidity chamber for 2 hours at 37°C. The list of antibodies is provided in Table S7.

Rolling circle amplification (RCA) and imaging: Probe ligation and labeling were performed using Duolink PLA detection-red kit (Sigma-Aldrich, DUO92008-100RXN) according to manufacturer’s instructions. RCA was performed using Duolink PLA detection-red kit (Sigma-Aldrich, DUO92008-100RXN) according to manufacturer’s instructions. Prior to each step, the cells were washed three times with 500 µL of wash buffer A (Sigma-Aldrich, DUO82036). To prepare for imaging, the cells were washed twice with wash buffer B (Sigma-Aldrich, DUO82036) and once with 1:100 wash buffer B (Sigma-Aldrich, DUO82036). Antifade mounting medium with DAPI (Sigma-Aldrich, DUO82040-5ML) was applied to each well. Coverslips (Corning®, CLS2980246) were placed onto the slides and sealed with a clear nail Top Coat. All imaging was performed using a Leica SP8 Confocal with Lightning Deconvolution Microscope, with a 60x objective. Images were processed using ImageJ. Statistical analyses were performed with GraphPad Prism 9.

### RIP-seq

Immunoprecipitation of RNA-protein complexes was conducted under native conditions utilizing either the anti-PHGDH antibody or IgG. Approximately 2x10^7^ HEK293T cells (ATCC, CRL3216) were lysed in 500 µL of the lysis buffer (50 mM Tris-HCl, pH 7.5 (Invitrogen™, 15567027), 100 mM NaCl (Thermo Fisher Scientific, AM9759), 1% Triton X-100 (Sigma-Aldrich, T8787-50ML), 0.1% SDS (Invitrogen™, AM9820), 0.5% Sodium Deoxycholate (Sigma-Aldrich, 30970-25G), and a protease inhibitor cocktail (Roche, 4693159001)) together with 200 U of RNasin® Plus Ribonuclease Inhibitor (40 U/µL, Promega, N2618) on ice for 30 minutes with occasional mixing and were then centrifuged at 15k rpm for 15 minutes. For each sample, 60 µL of Dynabeads™ Protein A (Invitrogen™, 10001D) were prepared in a 1.5 mL tube. The bead slurry was washed three times with 1 mL of the lysis buffer and resuspended in 100 µL of the lysis buffer. 5 µg of Rabbit anti-PHGDH antibody (Proteintech, 14719-1-AP) or Rabbit IgG isotype control (Abcam, AB37415) was mixed with the pre-washed beads at 4℃ for 2.5 hours on a rocking platform. Before use, the pre-equilibrated bead slurries were washed three times for 5 minutes with 1 mL of the lysis buffer.

Immunoprecipitation was conducted by incubating 500 µL of the cell lysate from each sample with the pre-equilibrated bead slurry as below at 4°C on a rocking platform overnight. Beads were sequentially washed twice with 1 mL of high salt buffer (50 mM Tris-HCl, pH 7.5 (Invitrogen™, 15567027), 1 M NaCl (Thermo Fisher Scientific, AM9759), 1 mM EDTA pH 8.0 (Invitrogen™, AM9260G), 1% Triton X-100 (Sigma-Aldrich, T8787-50ML), 0.1% SDS (Invitrogen™, AM9820), and 0.5% Sodium Deoxycholate (Sigma-Aldrich, 30970-25G)) and 1 mL of wash buffer (20 mM Tris-HCl, pH 7.5 (Invitrogen™, 15567027), 10 mM MgCl2 (Invitrogen™, AM9530G), and 0.2% Tween-20 (Sigma-Aldrich, P9416-100ML)). Complexes in each tube was released from beads by incubation with a mixing of 20 µL of Proteinase K (800 U/mL, NEB, P8107S) and 180 µL of PK buffer (50 mM Tris-HCl, pH 7.5 (Invitrogen™, 15567027)) and 10 mM MgCl2 (Invitrogen™, AM9530G)) at 50℃ for 40 minutes. The supernatant was collected and mixed with 1 mL of TRIzol Reagent (Invitrogen™, 15596026), and subsequently with 200 µL of chloroform (Acros Organics, A0425256), and was centrifuged at 14,000 rpm for 15 minutes at 4°C to extract RNA. The upper layer was collected. RNA was precipitated by the addition of 3 µL of glycogen (5 mg/mL, Invitrogen™, AM9510), 50% of 2-propanol (Sigma-Aldrich, I9516-500ML), and 10% of 3 M sodium acetate, pH 5.5 (Invitrogen™, AM9740) with an incubation at -80℃ overnight. The RNA was then pelleted by centrifugation at 14k rpm for 30 minutes at 4℃, washed with 1 mL of 75% ethanol (Sigma-Aldrich, 493546), and air-dried. The RNA was then suspended in 20 µL of UltraPure™ DNase/RNase-Free distilled water (Invitrogen™, 10977015).

A sequencing library was prepared by cDNA synthesis, amplification, fragmentation, and adaptor ligation using NEBNext® Low Input RNA Library Prep Kit (NEB, E6420) and sequenced by paired-end sequencing with 150 cycles from each end on an Illumina MiniSeq sequencer.

### PHGDH knockdown and overexpression

Knockdown in HEK293T: HEK293T cells (ATCC, CRL3216) were subcultured into a 6-well plate. The cells grew overnight, and the media were exchanged 6 hours before transfection. Cells were 70-90% confluent at the time of transfection. 100 pmol of 10 µM scrambled siRNA (Thermo Fisher Scientific, 4404021) (Control), PHGDH Silencer Select siRNA s514 (Thermo Fisher Scientific, s514, 5’-UAUUAGACGGUUAUUGCGTA-3’) (si-1) or s515 (Thermo Fisher Scientific, s515, 5’-UGAGCUCCAAGGUAAGAAGTG-3’) (si-2) was transfected to HEK293T cells with 10 µL of Lipofectamine 200 (Invitrogen™, 11668). Cells were harvested 48 h after transfection.

Knockdown in mNSC: mNSCs were subcultured into 6-well plates. After 24-hour culture, 200 pmol of 10 µM scrambled siRNA (Thermo Fisher Scientific, 4404021) (Control), Phgdh Silencer Select siRNA s108329 (Thermo Fisher Scientific, s108329, 5’-UUCAUCGAAGCUGUUGCCUGG-3’) (si-1) and s108330 (Thermo Fisher Scientific, s108330, 5’-UACUCGCACACCUUUCUUGCA-3’) (si-2) were transfected into mNSCs in each well of 6-well plates with 10 µL of Lipofectamine 200 (Invitrogen™, 11668). Cells were harvested 48 h after transfection.

Overexpression: WT and ED PHGDH were overexpressed in HEK293T (ATCC, CRL3216) cells by transfecting WT and ED PHGDH expression vectors, as well as an empty vector control. Cells were harvested 48 h after transfection,

### RT-qPCR

Total RNA was extracted from HEK293T cells and mNSCs using TRIzol reagent (Sigma, 93289) and further purified with chloroform. The RNA was then converted to complementary DNA (cDNA) using the SuperScript First-Strand Synthesis System (Thermo Fisher, 11904018) with random hexamers according to the manufacturer’s instructions. RT-qPCR was conducted with the Applied Biosystems PowerUp SYBR Green Master Mix (Thermo Fisher, A25742) to amplify the cDNA. Gene expression levels were quantified using the ΔΔCT method. TUBB mRNA and GAPDH mRNA were used as normalization controls for HEK293T cells and mNSCs, respectively.

For the RIP-qPCR assay, RNA was precipitated using anti-PHGDH antibody (Proteintech, 14719-1-AP) or IgG (Abcam, AB37415). Subsequently, the RNA underwent reverse transcription, and the RNA levels of BECN1, ATF4, and GAPDH were quantified using RT-qPCR, according to the previously described protocol. Quantification was performed using the ΔΔCT method, with GAPDH serving as the normalization control. The levels of BECN1 and ATF4 were normalized to those of GAPDH.

### Western blot

Cell pellets were suspended and lysed in 150 µL RIPA (Millipore, 20-188) containing protease inhibitor (cOmplete, Mini, EDTA-free Protease Inhibitor Cocktail, Roche, 11836170001) on ice for 30 minutes with occasional mixing and were then centrifuged at 14k rpm for 20 minutes. The supernatant was collected, and the protein concentrations were determined by Qubit Protein Broad Range (BR) Assay Kit (Invitrogen™, Q33211). 10% v/v β-mercaptoethanol (Gibco™, 21985023) was added to the 4X XT Sample Buffer (Bio-Rad, 1610791). Proteins from each sample were mixed with the 4X protein loading buffer to dilute to 1X and incubated at 100°C for 5 minutes. 30 μg of total proteins were resolved on a Tris-glycine 4-20% precast polyacrylamide gradient gel (Invitrogen™, XP00102BOX), together with a PageRuler™ Plus Prestained Protein Ladder (Thermo Scientific™, 26619) with 1X Tris/glycine/SDS running buffer (Bio-Rad, 1610772). SDS-PAGE was performed at 80 V for 45-60 minutes. For Western blotting, proteins were transferred onto an Immuno-Blot nitrocellulose membrane using an iBlot™ Gel Transfer Stack (Invitrogen™, IB301002), blocked with 5% milk in 1X TBST (Thermo Scientific™, 28360), and then incubated with diluted primary antibodies in the blocking solution at 4℃ overnight. The membrane was washed three times with 1x TBST and incubated with anti-Rabbit IgG HRP (Cell signaling, 7074) or anti-Mouse IgG HRP (Cell signaling, 7076) at a 1:1000 dilution in the blocking solution for 1 hour at room temperature. 1 mL of ECL Western Blotting Substrate solution (Thermo Scientific™, 32109) was prepared and uniformly spread on the membrane for imaging by Azure Imager c400 (Azure Biosystems) after TBST washes. The antibodies and dilutions used are shown in Table S7.

### Immunofluorescence

Cells were plated on the coverslips (Corning®, 354087) and cultured overnight. On the next day, cells were fixed with 4% formaldehyde (Thermo Fisher Scientific, 28906). For activated Caspase 3 (aCaspase3) and Nestin staining, cells were incubated with the block buffer (PBS containing 3% goat serum and 0.1% Triton X-100) at room temperature for 1 hour. For BrdU staining, cells were cultured with a medium containing 5 µM BrdU for 6 hours, washed three times with 1x PBS pH 7.2 (Life Technologies, 20012027), followed by a fixation with 4% formaldehyde (Thermo Fisher Scientific, 28906) in 1x PBS pH 7.2 (Life Technologies, 20012027) at room temperature for 30 minutes. The fixed cells were treated with 1 M HCl at 37 °C for 30 minutes, followed by incubation in a block buffer at room temperature for 1 hour.

Primary and secondary antibodies were diluted in antibody dilution buffer (PBS containing 3% goat serum and 0.1% Triton X-100). Samples were incubated with primary antibodies overnight at 4°C. The following day, samples were incubated with DAPI and fluorophore-conjugated secondary antibodies for 1 hour at room temperature. Finally, the slides were mounted for further analysis. The primary antibodies used in the immunostaining included: anti-activated Caspase-3 (CST, 9661S), anti-Nestin (BD Pharmingen™, 556309) and anti-BrdU (BD Biosciences, 560810). The secondary antibodies used in the immunostaining included: AlexaFluor488 goat anti-rabbit (Thermo FIsher, A11008) and AlexaFluor568 goat anti-mouse (Thermo FIsher, A11004).

For autophagosome staining, the Autophagy Assay Kit (AAT Bioquest, 23002) was used following the standard protocol provided by the manufacturer. A mixture of 2 µL of Component A and 1 mL of Component B was prepared, and fixed cells were incubated at 37°C for 30 minutes with the Component A/B mixture. After washing three times with Component C and air-drying, the coverslips were mounted onto slides with the antifade mounting medium containing DAPI (VECTASHIELD, H-1500) for further analysis.

### Quantification and statistical analysis Processing PRIM-seq read pairs

The following data processing steps are implemented in PRIMseqTools: https://github.com/Zhong-Lab-UCSD/PRIMseqTools. The raw sequencing read pairs were provided to Cutadapt 2.5 ^73^ to remove the 3′ linker sequence and the 5′ adaptor sequence. The remaining read pairs were subsequently subjected to Fastp 0.20.0 ^74^ and Python script to remove low-quality reads (average quality per base < Q20) and short reads (< 20 bp). The remaining read pairs were mapped to RefSeq transcripts ^75^ (based on GRCh38.p13, NCBI *Homo sapiens* Annotation Release 109.20190607) using BWA-MEM 0.7.12-r1039 ^76^ with the default parameters. The read pairs with one end mapped to the sense strand of a gene and the other end mapped to the antisense strand of a protein coding gene were identified as chimeric read pairs. Any duplicated chimeric read pairs were removed to obtain non-duplicate chimeric read pairs. A Chi-square test was carried out on every gene pair to test for RNA-protein association. The null hypothesis is that the mapping of one end of a chimeric read pair to a gene is independent of the mapping of the other end of this chimeric read pair to the other gene. The contingency table of this association test is given in Figure S2b. False discovery rate (FDR) was computed from the Benjamini-Hochberg procedure to control for family-wise errors in multiple testing.

### Downloading RNA-protein pairs from RNAInter database

RNA-protein associations were downloaded from the RNAInter database at http://www.rnainter.org/download/, specifying ‘Homo sapiens’ in both the ‘Species 1’’ column and the ‘Species 2’ column. iCLIP and HITS-CLIP derived RNA-protein associations were also downloaded from RNAInter by specifying ‘Homo sapiens’ in both the ‘Species 1’ column and the ‘‘Species 2’ column, ‘RBP’ in the ‘Category 2’ column, and the assay name (iCLIP or HITS-CLIP) in either the ‘strong’ column or the ‘weak’ column.

### Odds ratio calculation

The odds ratio was used to quantify the degree of overlap between two sets of RNA-protein associations (RPAs). The odds ratio (OR) of the following contingency table is calculated as OR = (A × D)/(C × B), where A, B, C, D are numbers of RNA-protein pairs in the corresponding cell in the contingency table.

|  | Within set II | Outside set II |
| --- | --- | --- |
| Within set I | A | B |
| Outside set I | C | D |

### GO term-defined subnetworks

The subnetwork associated with a GO term ^77^ was retrieved by the HuRPA proteins that were annotated by this GO term and all the edges connected with these proteins. GO term enrichment analysis was based on Chi-square tests. FDR was computed from the Benjamini-Hochberg procedure was used to control for family-wise errors. The protein classes belonging to the RNA processing proteins are categorized based on PANTHER.db ^78^. The entire HuRPA network was plotted with Gephi (0.9.2, https://gephi.org/) ^79^. All other network figures were plotted with Cytoscape ^80^.

### RNA-binding domains (RBDs) and RBD-binding motifs

RBDs were downloaded from RBPDB ^52^. The RBD classes with more than 100 domains captured by either RBDmap or pCLAP were used in this analysis. The Homer2 *de novo* module in Homer v5.0 ^81^ was applied to the RNA-end reads that are linked to the RRM class of RBDs to identify RBD-binding motifs, using all the RNA-end reads from the entire HuRPA as the background. The threshold for calling RBD-binding motifs was BH-corrected p-value smaller than 0.05 and more than 5% of the input sequences containing the motif.

### RIP-seq data analysis

The RIP-seq read pairs were mapped to RefSeq transcripts ^75^ (GRCh38.p13, NCBI *Homo sapiens* Annotation Release 109.20190607) using STAR 2.5.4b ^82^. FeatureCounts in Subread 2.0.6 ^83^ was applied to the resulting bam file to obtain the reads per million (RPM) for each gene. The PHGDH-associated RNAs were identified as the genes with BH-corrected p-values smaller than 0.05 (PHGDH vs. IgG, t test, two-sided) and an average RPM in the PHGDH libraries greater than 500.

### Precision and recall for RNA-protein association pairs

Precision and recall of HuRPA RNA-protein pairs were derived by comparing HuRPA RPAs with a reference set. Three reference sets were used, which are the RPAs in the RNAInter database (RNAInter RPAs), iCLIP-identified RPAs (iCLIP RPAs), HITS-CLIP identified RPAs (HITS-CLIP RPAs). An HuRPA RPA was considered matching a reference RPA only when both the RNA and the protein matched. The search space for precision-recall calculation was defined as the all the possible RNA-protein pairs between the HuRPA RNAs (7,248) and the proteins shared by HuRPA and the reference set.

## Data Availability

All PRIM-seq sequencing data has been deposited in GEO (GSE270010). All RIP-seq sequencing data has been deposited in GEO (GSE270009).

## Code Availability

PRIMseqTools and its source code and complete documentation are available at https://github.com/Zhong-Lab-UCSD/PRIMseqTools. A web interface for downloading, searching, and visualizing the HuRPA network is available at: https://genemo.ucsd.edu/prim.

## Author contributions

Z.Q., S.X., K.J., S.Z. designed the PRIM-seq technology, S.X. and K.J. generated the PRIM-seq libraries. Z.Q., X.W. carried out the data analysis. S.X. carried out the RNA-PLA experiments. J.C., W.Z. carried out RIP-seq and PHGDH perturbation experiments. Z.Q., S.X., and S.Z. took the lead in writing the manuscript. Z.Q., S.X., J.C., W.Z., J.L.C.R., S.Z. contributed to the interpretation of the results, provided critical feedback, and helped to shape the research, analysis, and manuscript.

