## Supplementary figures and tables for "Genome-Wide Mapping of RNA-Protein Associations via Sequencing"

Figure S1. PRIM-seq's experimental modules. (a) SMART-display. (b-i) REILIS (Reverse transcription, Incubation, Ligation, and Sequencing).

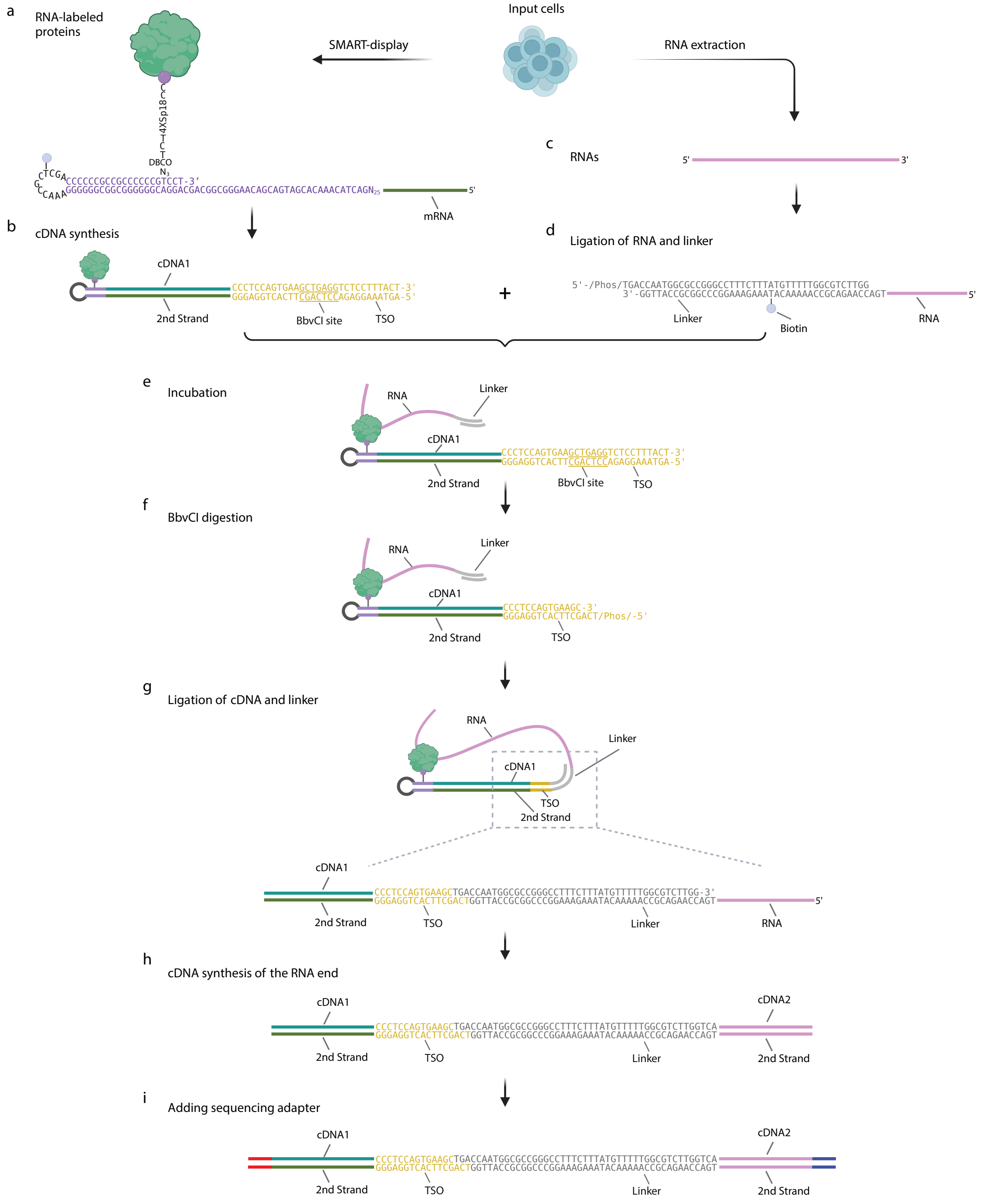

Figure S2. PRIM-seq data processing. (a) A cartoon showing the decoding of the protein-end and the RNA-end reads. As the sequencing reads from both ends are always from 5’ to 3’, the DNA-end reads are always antisense sequences and the RNA-end reads are alway sense strand sentences. (b) The contingency table for testing the independence of a RNA (RNA A) and a protein (Protein B) from PRIM-seq data. Xij are the read counts. A Chi-square test is constructed from this contingency table for each pair of RNA and protein. (c) Flowchart of PRIMseqTools for processing PRIM-seq data. Adaptor sequences were trimmed (Adaptor trimming) and low quality reads were removed (Quality filtering). The resulting read pairs were mapped to Refseq genes (Mapping). The read pairs with the two ends mapped to two different genes are retained (Identification of chimeric read pairs) and deduplicated. Non-duplicated chimeric read pairs with one end sensely mapped to a gene and the other end antisensely mapped to a protein-coding gene (RNA/protein end assignment) were used as the input for the Chi-square test (Statistical test).

a

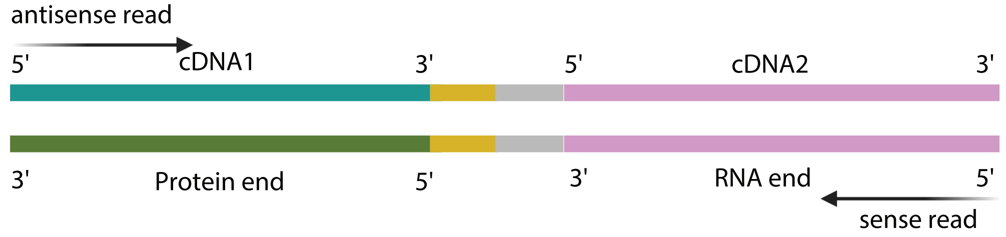


b

Contingency table

| | | | | |
|---|---|---|---|---|
| | | **Mapped to Protein B** | | |
| | | No | Yes | |
| **Mapped to RNA A** | No | $X_{00}$ | $X_{01}$ | |
| | Yes | $X_{10}$ | $X_{11}$ | |
| | | | | Total number of deduplicated valid read pairs |

c

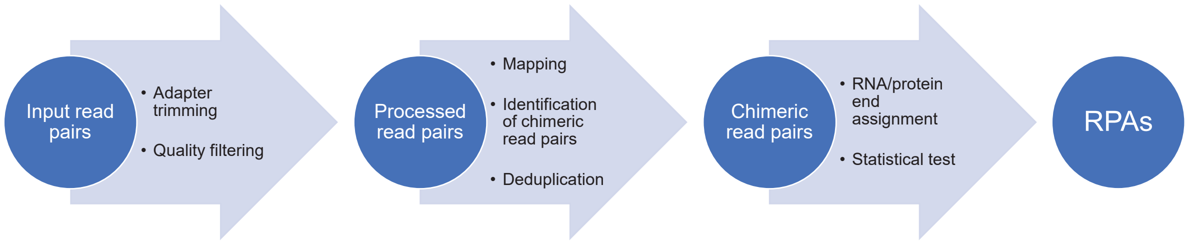

Figure S3. Reproducibility of PRIM-seq libraries. (a,c,e,g) Venn diagrams of the identified RNA-protein pairs between any two of the three HEK293T replicates (HEK-1, HEK-2, HEK-3) (a,c,e) and between the two K562 replicates (K562-1, K562-2) (g). (b,d,f,h). The odds ratio, i.e. degree of overlap (y axis) increases as the threshold for calling the RNA-protein associations (RPAs) from each replicate (x axis) increases. X: the expected number of read pairs from a randomly chosen RNA-protein pair. nX: n multiplies X.

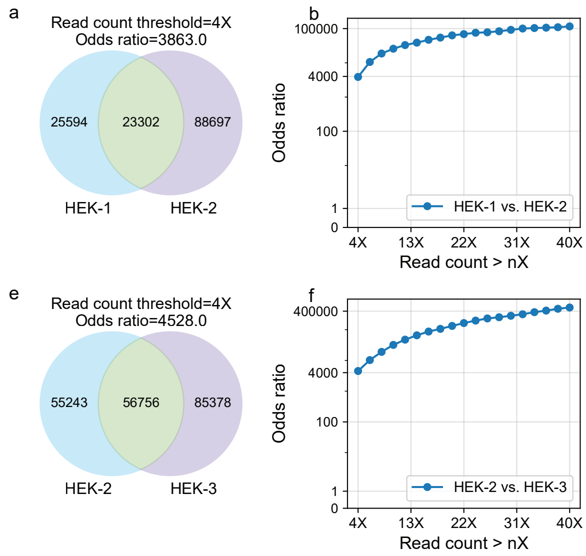

Figure S4. HuRPA RNAs and HuRPA proteins. (a-b) Distribution of HuRPA RNAs by RNA types, counted by the number of RNA genes (a) and the number of RNA-protein pairs (b) in HuRPA. (c) Upset plot of the intersections of RBPs of databases: RBP2GO, RBPDB, ATtRACT, hRBPome, RBPbase, and starBase. (d) Venn diagram showing significant overlaps of Undatabased HuRPA proteins with pCLAP captured proteins and RBDmap captured proteins. P-values are derived from Chi-square tests. (e) Degree rank distributions of Undatabased HuRPA proteins captured by RBDmap, by pCLAP, and other human proteins in HuRNA-protein association. Lower rank indicates higher degrees in HuRNA-protein association. (e) Rank distribution of the degrees of Undatabased HuRPA proteins, categorized by pCLAP captured proteins (blue), RBDmap (orange) captured proteins, and the rest (green). The protein with most associated RNAs (highest degree) is ranked on top (rank = 1). pCLAP and RBDmap captured proteins exhibited larger degrees than the other Undatabased HuRPA proteins.

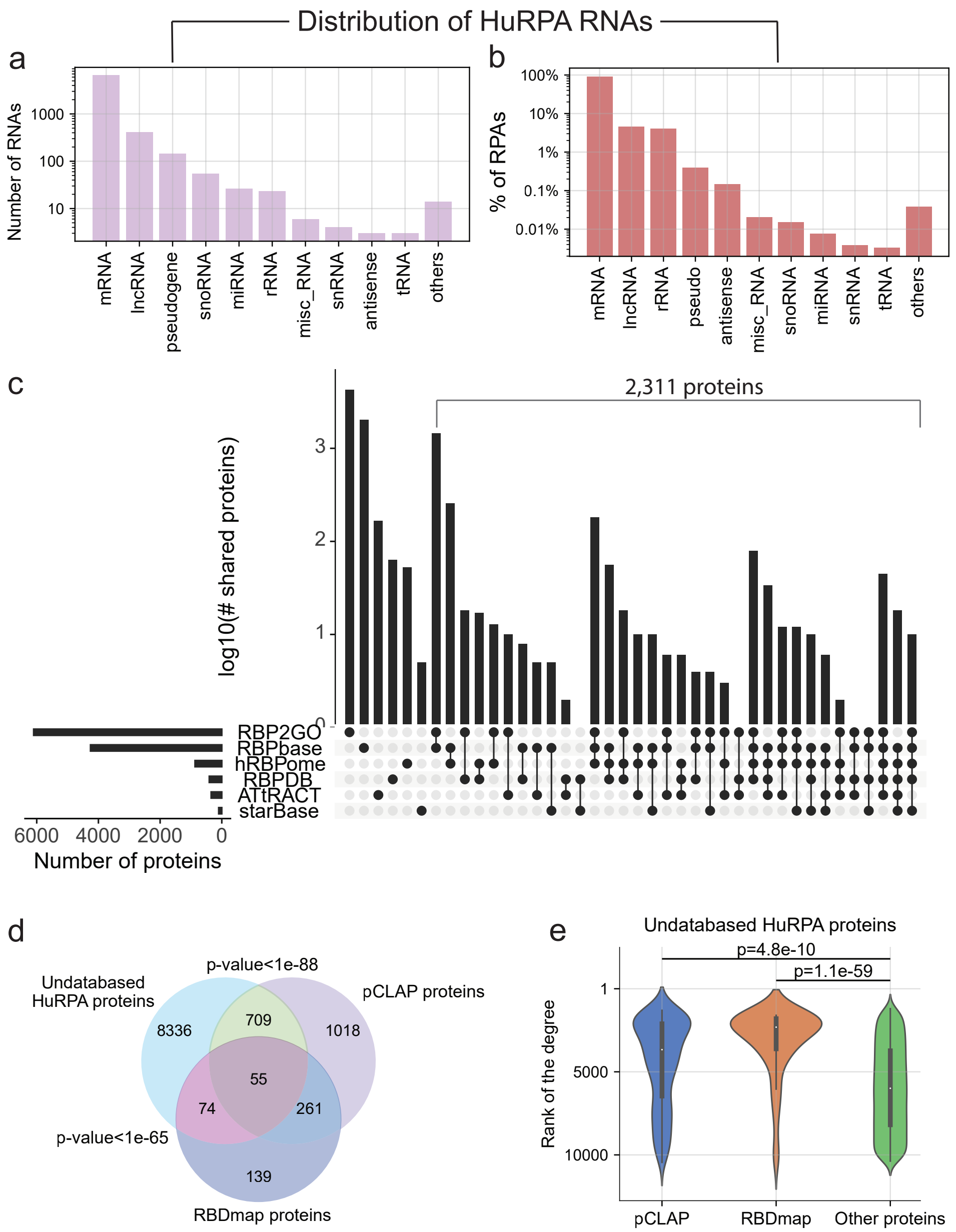

Figure S5. Gene Ontology analysis. (a,d,f) The number of genes (x axis) in each GO term (dot) is plotted against the significance level (y axis) of this term in the HuRPA proteins. To avoid very general GO terms, we restricted our analysis to Biological Process (BP) terms with no more than 1,000 genes per term (a), and Cellular Component (CC) and Molecular Function (MF) terms with no more than 100 genes per term (d, f). (b,e,g) The numbers of RNA-protein associations (RPAs), RNAs, and proteins in the HuRPA subnetworks associated with the most significant GO terms: “RNA processing” (b), “cytoplasmic stress granule” (e), and “translation factor activity, RNA binding” (g). (c) Distribution of "RNA processing" associated HuRPA proteins across different protein classes.

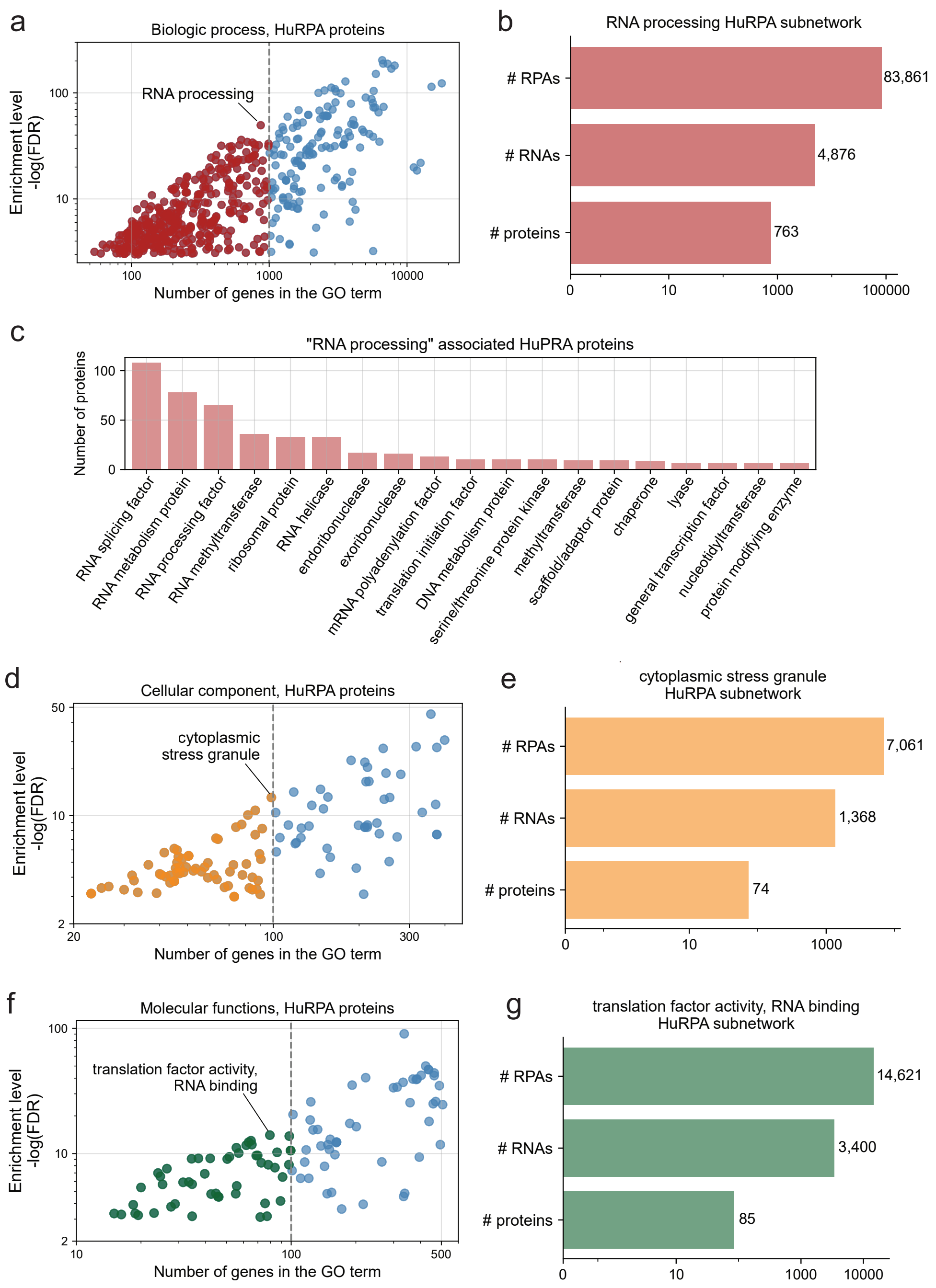

Figure S6. Precision and recall of RNA-protein associations. (a-c) Precision (y axis) and recall (x axis) are derived from comparing the RNA-protein pairs in HuRPA (rightmost black dot) with three reference sets: the RNA-protein pairs in the entire RNAInter database (a), iCLIP-identified RNA-protein pairs (b), and HITS-CLIP-identified RNA-protein pairs (c). Increasing the threshold for calling RNA-protein associations (arrow) from PRIM-seq data resulted in higher precision and lower recall (black dots). Randomly sampled RNA-protein pairs resulted in significantly smaller precisions and recalls (gray dots). (d-f) Precisions and recalls of the highly connected HuRPA subnetworks. The subnetworks are derived from retaining the HuRPA proteins with their degrees ranked among the top 10% (green), 25% (pink), 50% (blue), and 100% (black) in HuRPA. The HuRPA subnetworks with higher degrees (more connections) exhibited higher precisions and recalls.

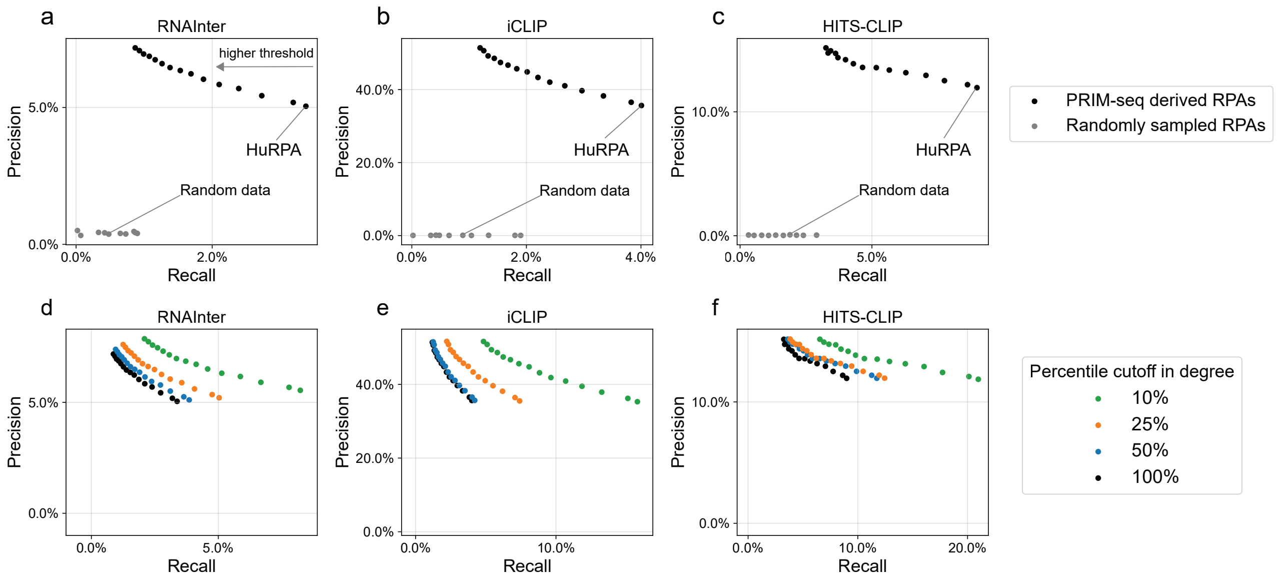

Figure S7. RNA binding domains (RBDs) and RNA-binding motifs. (a) The probability density function (blue curve) of the number of protein-end reads that can overlap with RBDs as a random variable, compared to the observed number of protein-end reads from PRIM-seq that overlapped with RBDs (vertical line). The p-value is an empirical p-value. (b) The RBDs (red blocks) on the HNRNPR protein sequence (upper panel) co-localize with the peaks of PRIM-seq's protein-end reads (lower panel). (c) Proportions of RBPs and RBDs captured by HuRPA, categorized by RBD classes. (d) The enrichment level (-log10(p-value, x axis) of each type of RBDs vs. the number of RBDs captured in HuRPA proteins (y axis) for each RBD class. (e) Comparison of known consensus RNA sequences of the RBD-binding motifs (blue) of the RRM RBD with the de novo motifs identified from the RNA-end reads (multi-color logo) that are linked to the RRM-mapped protein-end reads.

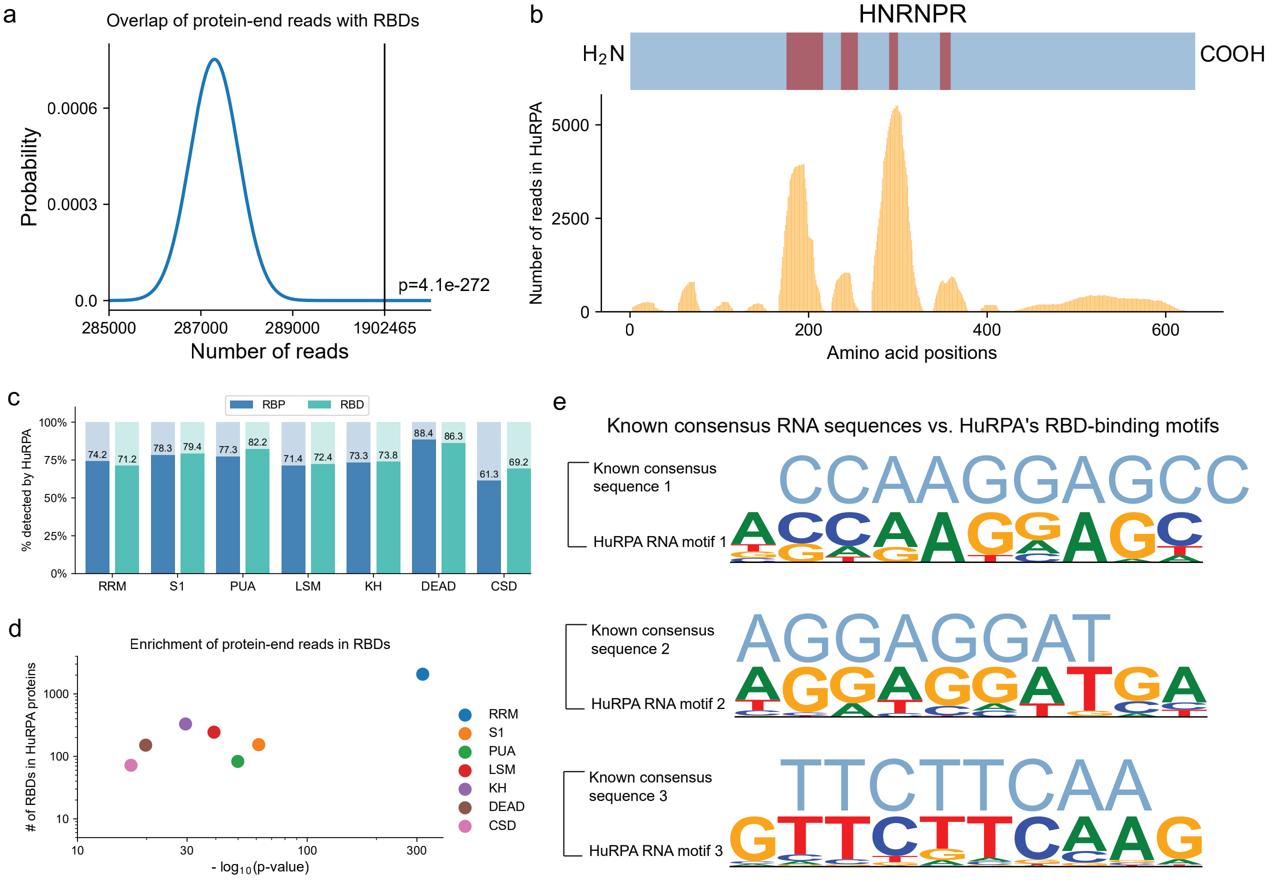

Figure S8. Reproducing a previously published RNA-PLA experiment as a sanity check. (a-b) Representative images of RNA-PLA signals (red) with Smith antibody and with (a) and without the U1 RNA probe (b). Nuclei are DAPI stained (blue). Scale bars = 20 μm. (c) The average number of RNA-PLA foci per cell (y axis) in each group. # of cells: the number of cells analyzed. Error bar: SEM. P-value is derived from a two-sided t test.

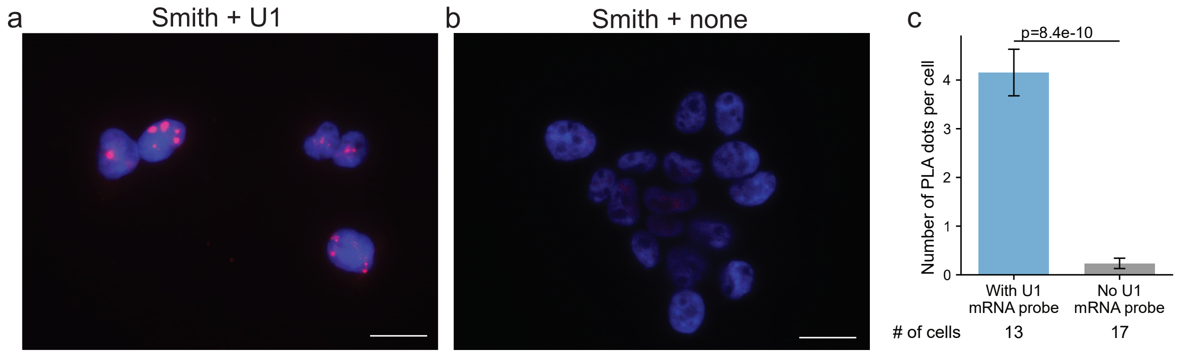

Figure S9. RNA-PLA analysis of RNA-protein pairs involving LINC00339 noncoding RNA. (a-c) Representative images of the tests (protein name + LINC00039) (a), antibody-only controls (protein name + none) (b), other negative controls including no-probe-no-antibody control (none + none), RNA probe-only control (none + LINC00339), and four RNA-protein pairs not included in HuRPA (GFP + LINC0039, CD40 + LINC0039, CD32 + LINC0039, LTBR + LINC0039 (c). Quantification is provided in Figure 3k. Blue: DAPI staining. Red: RNA-PLA signal. Scale bar = 20 μm.

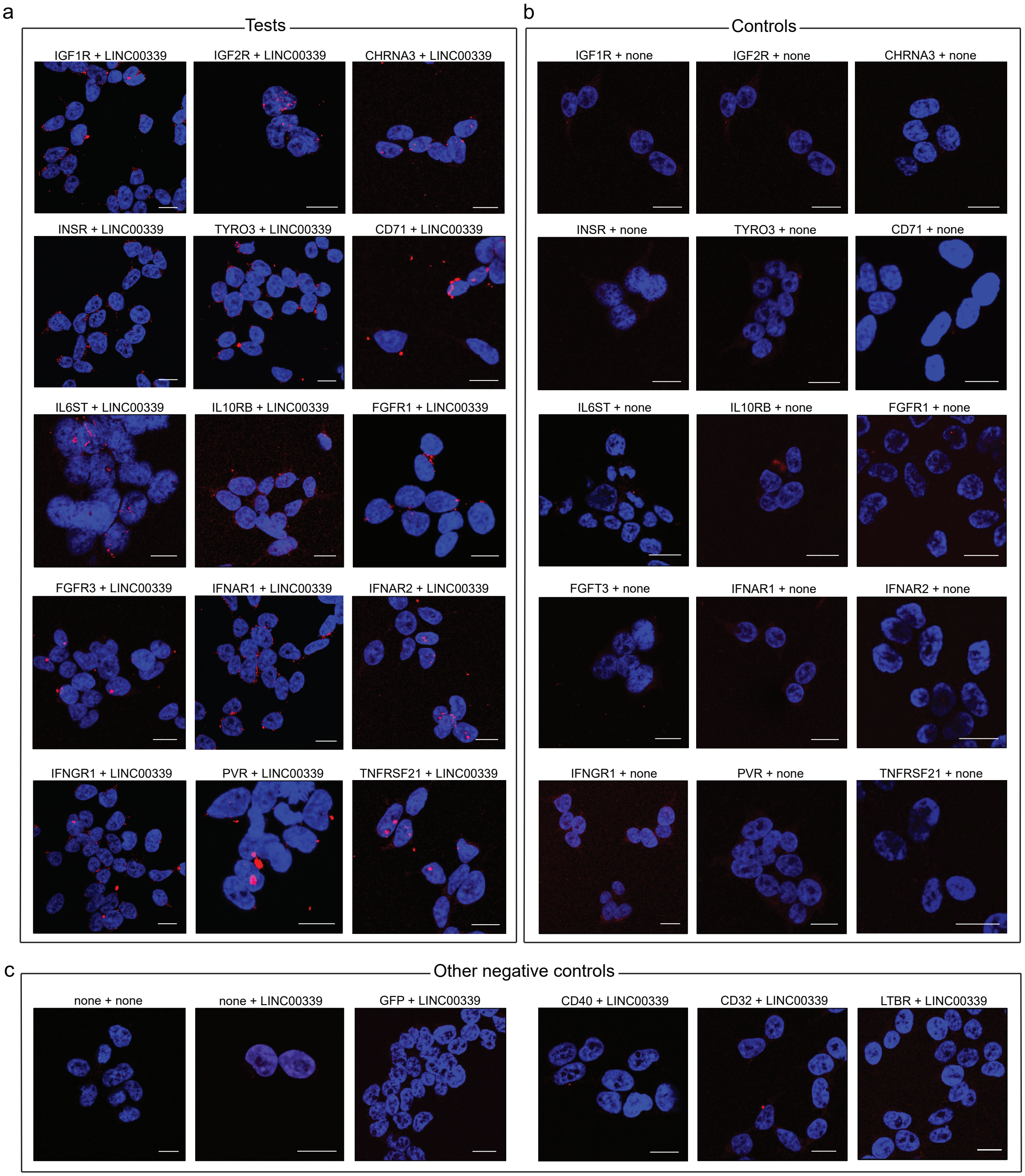

Figure S10. RIP-seq and RNA-PLA analysis of PHGDH. (a) Categorization of PHGDH-associated RNAs in HuRPA by RNA classes. (b) The average RPM (reads per million, y axis) of each RNA gene (dot) in PHGDH RIP-seq vs. the enrichment level (-log10(adjusted p-value), x axis) of PHGDH as compared to IgG. Purple and red dots: RIP-seq identified PHGDH-associated RNAs. Red dots: PHGDH-associated RNAs identified by both RIP-seq and HuRPA. (c) Comparison of PHGDH-associated RNAs in HuRPA and detected by RIP-seq. Odds ratio (y axis) is greater than 1, indicating a significant overlap. Error bar: 95% confidence interval. As the threshold for calling PHGDH-associated RNAs from RIP-seq (x axis) increases, the odds ratio also increases, indicating a stronger overlap. (d) Among the PHGDH-associated RNAs in HuRPA, the RNA-end reads (y axis) of the RNAs detected (blue) by PHGDH RIP-seq are more than the RNAs not-detected by PHGDH RIP-seq (orange). (e-h) Representative RNA-PLA images of the PHGDH protein and ATF4 mRNA (e), RNA-probe-only control (f), antibody-only control (g), and no-antibody-no-RNA-probe control (h). Scale bar = 20 μm.

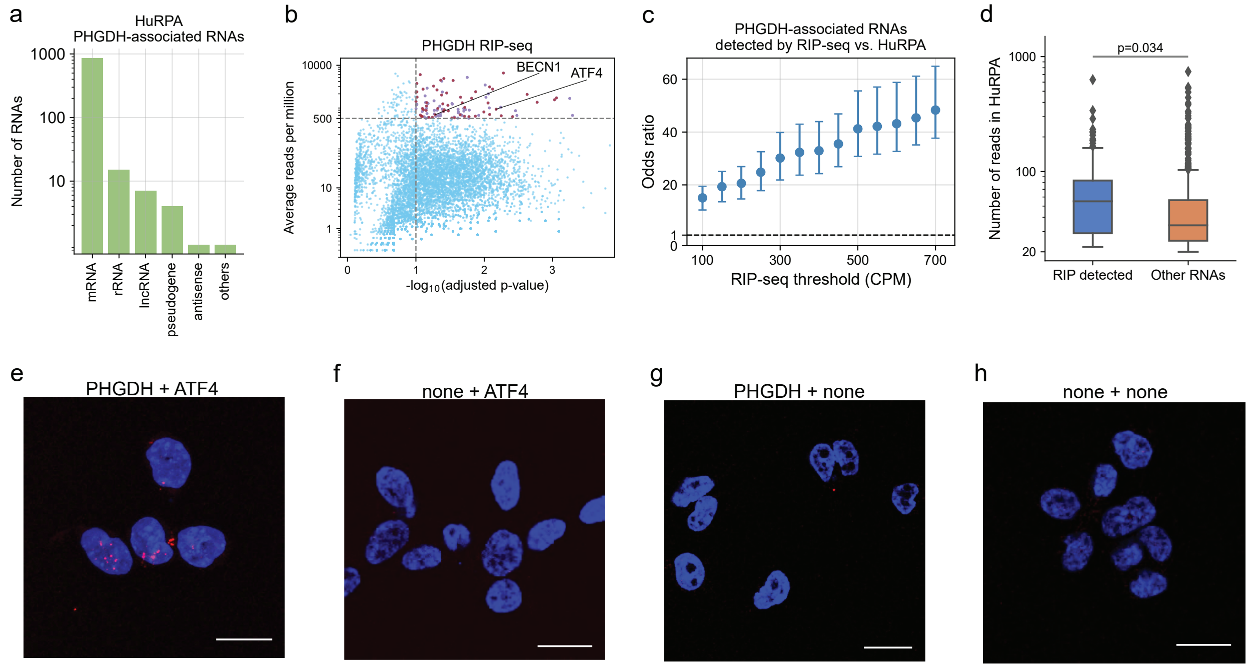

Figure S11. Cellular responses to PHGDH knockdown in HEK293T cells. Immunofluorescence staining and quantification of autophagosome (a,b), BrdU (c,d), and activated Caspase 3 (aCaspase3) (e,f) in scramble siRNA (Control) and PHGDH-targeting siRNAs (si-1, si-2) treated HEK293T cells. P-values are derived from two-sided t-tests. Error bar: SEM. n: number of replicates.

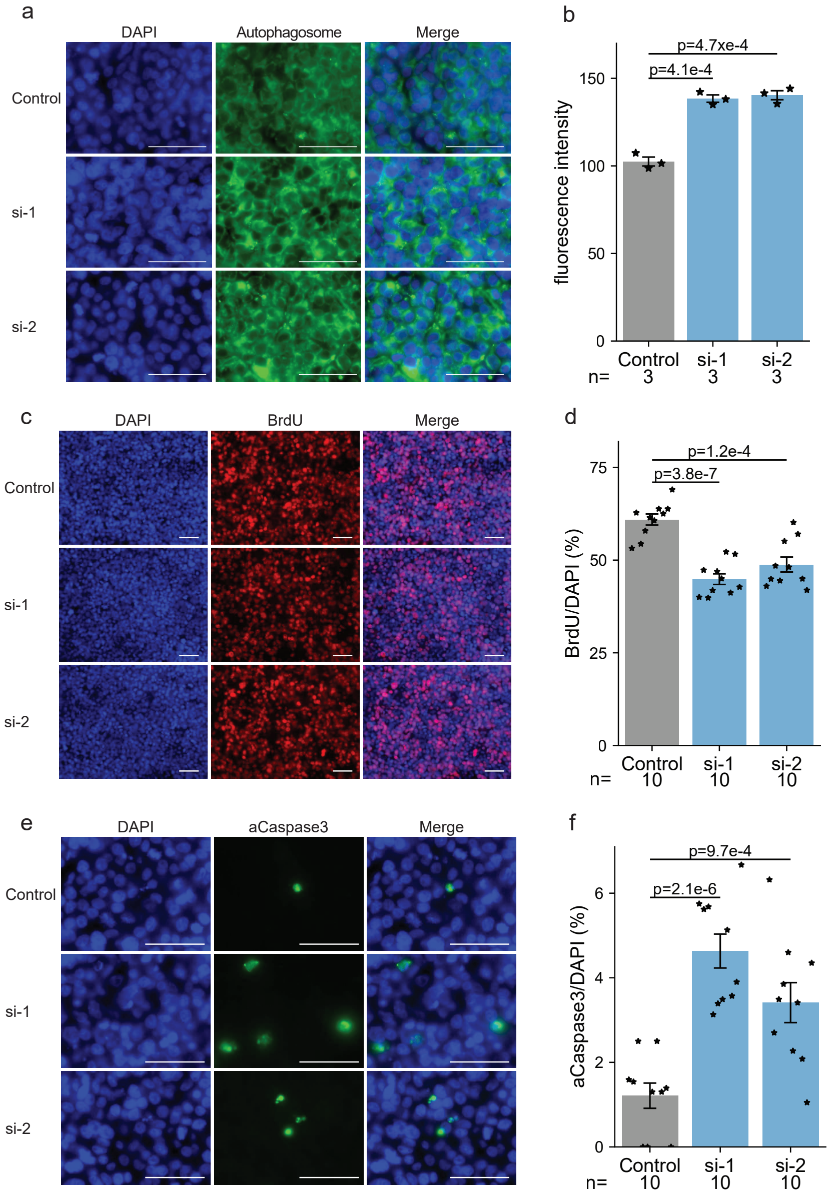

Figure S12. Cellular responses to PHGDH knockdown in mouse neural stem cells (mNSCs). P-values are derived from two-sided t-tests. Error bar: SEM. n: number of replicates. (a) PHGDH RNA levels in scramble siRNA (Control) and PHGDH-targeting siRNAs (si-1, si-2) treated mNSCs. (b-i) Immunofluorescence staining and quantification of autophagosome (b,c), BrdU (d,e), and activated Caspase 3 (aCaspase3) (f,g) in scramble siRNA (Control) and PHGDH-targeting siRNAs (si-1, si-2) treated mNSCs. (h-j) Cell morphology analysis. Immunofluorescence staining of Nestin (h), normalized average dendrite length (i), and the number of dendrite intersections (y axis) as a function of distance from the soma (x axis) (j) in scramble siRNA (Control) and PHGDH-targeting siRNAs (si-1, si-2) treated mNSCs.

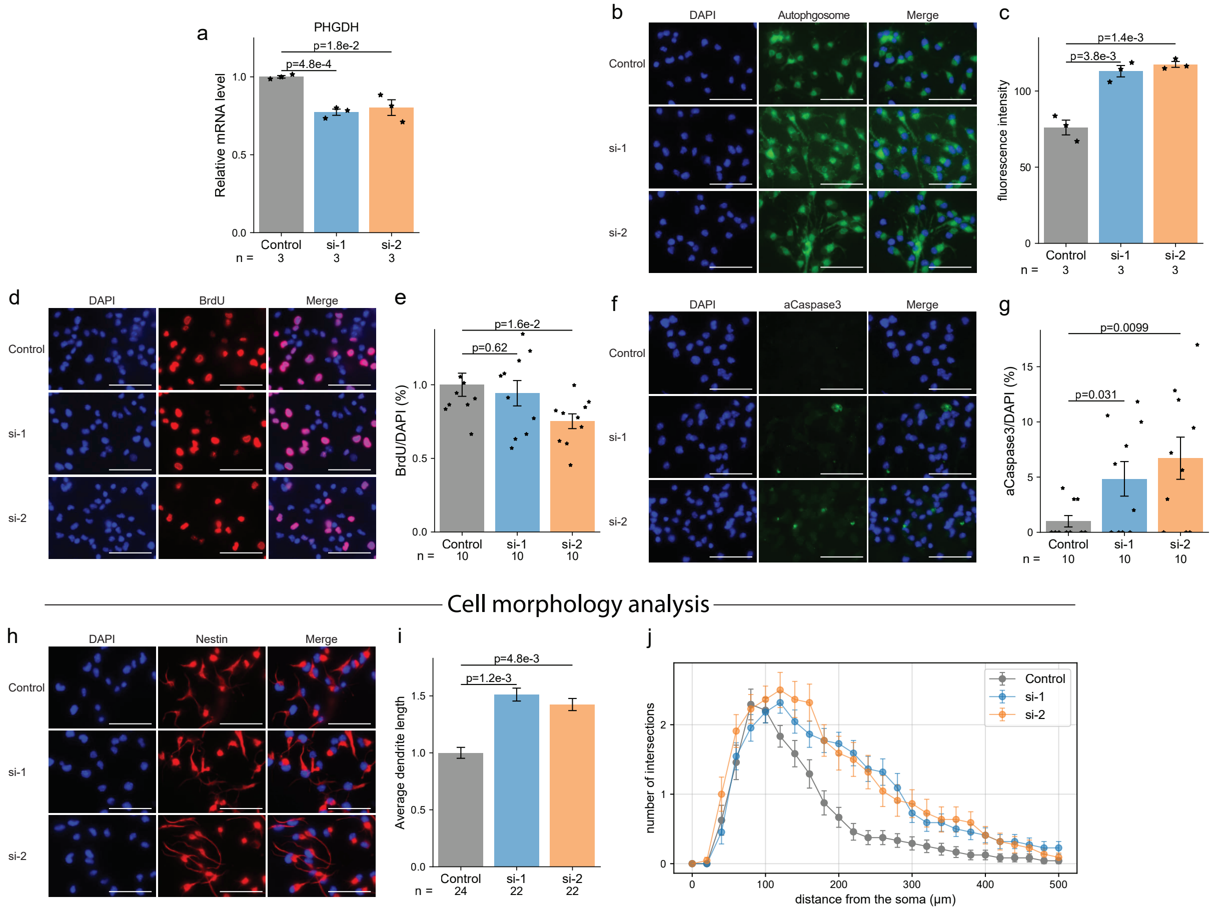

Figure S13. Illustration of RNA-linker ligation.

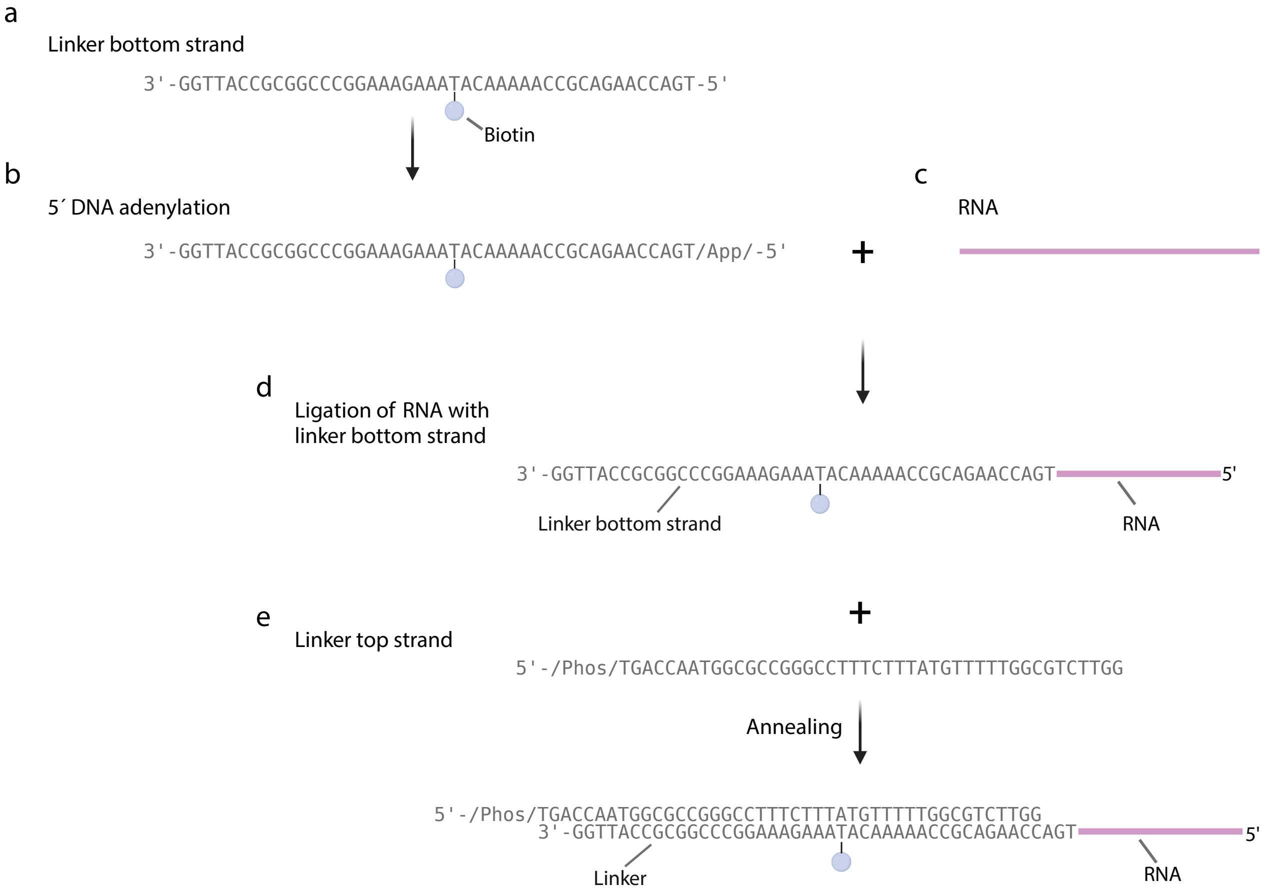

Table S1. Summary of PRIM-seq libraries. The PRIM-seq libraries are indexed by Library ID and Experiment ID (Exp ID), annotated with the cell line used (Cell line), the total number of read pairs (# read pairs), the number of read pairs mapped to the transcriptome (# mapped read pairs), and the number of non-duplicate read pairs with one end mapped to the sense strand of a gene and the other end mapped the antisense strand of a protein coding gene (# non-duplicate chimeric read pairs).

| Library ID | Exp ID | Cell line | # read pairs | # mapped read pairs | # non-duplicated chimeric read pairs |
|---|---|---|---|---|---|
| HEK-1 | 1 | HEK293T | 409,132,179 | 188,352,224 | 5,814,154 |
| HEK-2 | 2 | HEK293T | 656,928,590 | 400,343,720 | 14,096,530 |
| HEK-3 | 3 | HEK293T | 797,158,755 | 420,122,030 | 20,246,532 |
| K562-1 | 4 | K562 | 717,735,040 | 366,313,999 | 12,794,460 |
| K562-2 | 5 | K562 | 532,026,785 | 360,332,965 | 9,425,554 |

Table S2. Public datasets used in this work. The previously published datasets are indexed by their names (Name). The RNA-protein association data (RNAInter, eCLIP, and HITS-CLIP) are annotated with the number of RNA-protein associations (# RPAs), the number of RNAs (# RNAs), and the number of proteins (# proteins) in each dataset (row). The RBP screening data (pCLAP and RBDmap) are annotated with the number of RNA-binding proteins (# RBPs) and the number of RNA-binding domains (# RBDs).

| **Name** | **Description** | **# RPAs** | **# RNAs** | **# proteins** |
|---|---|---|---|---|
| **RNAInter** | All the experimentally-derived human RNA-protein interactions in RNAInter, downloaded from http://www.rnainter.org/download/ | 816,996 | 23,674 | 8,164 |
| **HITS-CLIP** | high-throughput sequencing of RNA isolated by crosslinking immunoprecipitation detected human RNA-protein interactions that are included in RNAInter | 62,310 | 14,184 | 39 |
| **iCLIP** | Individual-nucleotide resolution UV crosslinking and immunoprecipitation detected human RNA-protein interactions that are included in RNAInter | 128,030 | 15,669 | 33 |
| | | | **# RBPs** | **# RBDs** |
| **pCLAP** | Peptide crosslinking and affinity purification detected RBPs and RBDs, by Mullari, Lyon, Jensen, & Nielsen, 2017 | | 2,043 | 4,751 |
| **RBDmap** | RBDmap detected RBPs and RBDs, Castello, by Fischer et al. 2016 | | 529 | 1,611 |

Table S3. RNA-binding domains (RBDs) in HuRPA proteins. Each RBD class (row) is annotated with the number of HuRPA proteins harboring this class of RBDs (# proteins in HuRPA), the number of RBDs harbored in these HuRPA proteins (# domains in HuRPA), the number of protein-end reads aligned these RBDs (# protein-end reads), and the p-values derived from one-sided binomial tests (p-value).

| RBD class | # proteins in HuRPA | # domains in HuRPA | # protein-end reads | p-value |
|---|---|---|---|---|
| **RRM** | 861 | 2,067 | 1,049,071 | $8.6 \times 10^{-319}$ |
| **LSm** | 105 | 244 | 74,130 | $5.3 \times 10^{-40}$ |
| **S1** | 65 | 154 | 141,318 | $3.2 \times 10^{-62}$ |
| **KH** | 129 | 330 | 68,579 | $2.9 \times 10^{-30}$ |
| **PUA** | 34 | 83 | 96,848 | $1.4 \times 10^{-50}$ |
| **DEAD** | 38 | 151 | 35,772 | $1.5 \times 10^{-20}$ |
| **CSD** | 19 | 72 | 10,610 | $8.0 \times 10^{-18}$ |

Table S4. Proteins tested with LINC00339 in the RNA-PLA assay. Each protein (row) is annotated with whether it is associated with LINC00339 in HuRPA (Test if yes, Control if no), and its subcellular localization as shown in the Human Protein Atla database.

| Protein | Test/Control | Subcellular localization |
|---|---|---|
| IGF1R | Test | Plasma membrane (predicted) |
| IGF2R | Test | Golgi apparatus and Vesicles |
| CHRNA3 | Test | Plasma membrane and Nuclear speckles |
| INSR | Test | Vesicles and Plasma membrane |
| TYRO3 | Test | Plasma membrane (predicted) |
| CD71 | Test | Endosomes and Lysosomes; and is predicted to be secreted |
| IL6ST | Test | Plasma membrane and Golgi apparatus; and is predicted to be secreted |
| IL10RB | Test | Cytosol |
| FGFR1 | Test | Plasma membrane (predicted) |
| FGFR3 | Test | Endoplasmic reticulum; Secreted (predicted) |
| IFNAR1 | Test | Plasma membrane (predicted) |
| IFNAR2 | Test | Plasma membrane (predicted); Secreted (predicted) |
| IFNGR1 | Test | Plasma membrane |
| PVR | Test | Nucleoplasm, Vesicles and Plasma membrane; and is predicted to be secreted |
| TNFRSF21 | Test | Plasma membrane and Cytosol |
| GFP | Control | N/A |
| CD40 | Control | Secreted (predicted) |
| CD32 | Control | Plasma membrane and Golgi apparatus |
| LTBR | Control | Golgi apparatus |

Table S5. Summary of RIP-seq libraries. Each library (row) is annotated with a Library ID and an an Experiment ID (Exp ID), the RIP antibody (Antibody), the cell line used for generating the library (Cell line), the total number of read pairs (# read pairs), and the number of uniquely mapped read pairs (# uniquely mapped read pairs).

| Library ID | Exp ID | Antibody | Cell line | # read pairs | # uniquely mapped read pairs |
|---|---|---|---|---|---|
| PHGDH-1 | 1 | anti-PHGDH | HEK293T | 5,493,909 | 4,724,012 |
| IgG-1 | 1 | IgG | HEK293T | 6,337,515 | 5,314,700 |
| PHGDH-2 | 2 | anti-PHGDH | HEK293T | 2,874,421 | 1,968,860 |
| IgG-2 | 2 | IgG | HEK293T | 3,111,112 | 2,501,481 |

Table S6. Antisense oligonucleotide RNA probes used in the RNA-PLA assay. Each probe (row) is annotated with a Probe ID, the target RNA, and its oligonucleotide sequence. Dig_N: Digoxigenin.

| **Probe ID** | **Target RNA** | **Oligonucleotide sequence** |
|---|---|---|
| U1 | U1 | 5’-CTGGGAAAACCACCTTCGTGATCATGGTATCTCCCCTGCC AGGTAAGTATAAAA /Dig_N/-3’ |
| L1 | LINC00339 | 5’-ATTTCTTTGTGTCTCTGGGTACACTTCAGTCTCAATTCTGG AACTCAAAA/Dig_N/-3 |
| L2 | LINC00339 | 5’-CTCATATGCAGGCTGAGCAAGAAGCAAGCAGCAAGACTT AGCCAGTAAAA/Dig_N/-3’ |
| L3 | LINC00339 | 5’-CAAGGGCACCCCAAATGAGTTACTGGTGGGGTTACATAA CCTTTCAAAAA/Dig_N/-3’ |
| A1 | ATF4 mRNA | 5’-ATTCGAAGGTGTCTTTGTCGGTTACAGCAACGCTGCTGC TGAATGCAAAA/Dig_N/-3’ |
| A2 | ATF4 mRNA | 5’-CAGCTCTAAACTAAAGGAATGATCTGGAGTGGAGGACAG GACCCCTAAAA/Dig_N/-3’ |
| A3 | ATF4 mRNA | 5’-GGAACACACAGCTACAGCACTCTATGTACAAGCACATTGA CGCTCCAAAA/Dig_N/-3’ |

Table S7. Antibodies used in this study. The antibodies are annotated by the target protein (Target), vendor and catalog number (Source), species and clonal information (Description), dilution used in this study (Dilution), and the assay that used this antibody (Assay).

| Target | Source | Description | Dilution | Assay |
|---|---|---|---|---|
| Smith | Sigma, MABF2793-100UL | Mouse monoclonal | 1:200 | RNA-PLA |
| Digoxin | Sigma, SAB4200669-100UL | Mouse monoclonal | 1:200 | RNA-PLA |
| IGF1R | Proteintech, 20254-1-AP | Rabbit polyclonal | 1:200 | RNA-PLA |
| IGF2R | Proteintech, 20253-1-AP | Rabbit polyclonal | 1:200 | RNA-PLA |
| CHRNA3 | Proteintech, 10333-1-AP | Rabbit polyclonal | 1:200 | RNA-PLA |
| INSR | Proteintech, 20433-1-AP | Rabbit polyclonal | 1:200 | RNA-PLA |
| TYRO3 | Proteintech, 28513-1-AP | Rabbit polyclonal | 1:200 | RNA-PLA |
| CD71 | Proteintech, 10084-2-AP | Rabbit polyclonal | 1:200 | RNA-PLA |
| IL6ST | Proteintech, 67766-1-Ig | Mouse monoclonal | 1:200 | RNA-PLA |
| IL10RB | Proteintech, 15102-1-AP | Rabbit polyclonal | 1:200 | RNA-PLA |
| FGFR1 | Proteintech, 60325-1-Ig | Mouse monoclonal | 1:200 | RNA-PLA |
| FGFR3 | Proteintech, 66954-1-Ig | Mouse monoclonal | 1:200 | RNA-PLA |
| IFNAR1 | Proteintech, 13083-1-AP | Rabbit polyclonal | 1:200 | RNA-PLA |
| IFNAR2 | Proteintech, 10522-1-AP | Rabbit polyclonal | 1:200 | RNA-PLA |
| IFNGR1 | Proteintech, 10808-1-AP | Rabbit polyclonal | 1:200 | RNA-PLA |
| PVR | Proteintech, 27486-1-AP | Rabbit polyclonal | 1:200 | RNA-PLA |
| TNFRSF21 | Proteintech, 66754-1-Ig | Mouse monoclonal | 1:200 | RNA-PLA |
| GFP | ORIGENE, TA150041 | Mouse monoclonal | 1:200 | RNA-PLA |
| CD40 | Proteintech, 28158-1-AP | Rabbit polyclonal | 1:200 | RNA-PLA |
| CD32 | Proteintech, 15625-1-AP | Rabbit polyclonal | 1:200 | RNA-PLA |
| LTBR | Proteintech, 20331-1-AP | Rabbit polyclonal | 1:200 | RNA-PLA |
| PHGDH | Proteintech, 14719-1-AP | Rabbit polyclonal | 1:1000 | Western Blot |
| BECN1 | Proteintech, 11306-1-AP | Rabbit polyclonal | 1:5000 | Western Blot |
| ATF4 | Proteintech, 10835-1-AP | Rabbit polyclonal | 1:1000 | Western Blot |
| BCLAF1 | Proteintech, 26809-1-AP | Rabbit polyclonal | 1:2000 | Western Blot |
| VCL | Proteintech, 66305-1-lg | Mouse monoclonal | 1:10000 | Western Blot |
| ATCB | Proteintech, 66009-1-lg | Mouse monoclonal | 1:20000 | Western Blot |
